# Buffering and total calcium levels determine the presence of oscillatory regimes in cardiac cells

**DOI:** 10.1101/2020.02.14.949180

**Authors:** M. Marchena, Blas Echebarria, Yohannes Shiferaw, Enrique Alvarez-Lacalle

## Abstract

Calcium oscillations and waves are often behind instances of extra depolarization in cardiac cells, eventually giving rise to life-threatening arrhythmias. In this work, we study the conditions for the appearance of calcium oscillations in both a detailed subcellular model of calcium dynamics and a minimal model that takes into account just the minimal ingredients of the calcium toolkit. To avoid the effects of homeostatic changes and the interaction with the action potential we consider the somewhat artificial condition of a cell without pacing and with no calcium exchange with the extracellular medium. This permits us to isolate the main reasons responsible for the oscillations by controlling externally the total calcium content of the cell. We find that as the calcium content is increased, the system transitions between two stationary states, corresponding to one with closed ryanodine receptors (RyR) and most calcium in the cell stored in the sarcoplasmic reticulum (SR), and another, with open RyRs and a depleted SR. In between these states, calcium oscillations may appear. This transition depends very sensitively in the amount of buffering in the cell. We find, for instance, that at high values of calsequestrin (CSQ) oscillations disappear, while they are present for a broad range of parameters at low values of CSQ. Using the minimal model, we can relate the stability of the oscillating state to the nullcline structure of the system, and find that its range of existence is bounded by a homoclinic and a Hopf bifurcation.

**Author summary:** In cardiac cells, calcium plays a very important role. An increase in calcium levels is the trigger used by the cell to initiate contraction. Besides, calcium modulates several transmembrane currents, affecting the cell transmembrane potential. Thus, dysregulations in calcium handling have been associated with the appearance of arrhythmias. Often, this dysregulation results in the appearance of periodic calcium waves or global oscillations, providing a pro-arrhythmic substrate. In this paper, we study the onset of calcium oscillations in cardiac cells using both a detailed subcellular model of calcium dynamics and a minimal model that takes into account just the minimal ingredients of the calcium toolkit. Both reproduce the main experimental results and link this behavior with the presence of different steady-state solutions and bifurcations that depend on the total amount of calcium in the cell and in the level of buffering present. We expect that this work will help to clarify the conditions under which calcium oscillations appear in cardiac myocytes and, therefore, will represent a step further in the understanding of the origin of cardiac arrhythmias.

## Introduction

Cardiovascular diseases represent one of the main causes of death worldwide [1]. Often, mortality is related to the appearance of rapid cardiac rhythms, such as tachycardia and fibrillation, that result in contractibility loss, reducing cardiac output and eventually leading to sudden cardiac death [2]. Although the onset of rapid arrhythmias can be due to a large variety of factors [3], including changes in the properties of cardiac tissue [4], often arrhythmias are triggered by spontaneous intracellular calcium releases [5, 6]. In cardiac cells, calcium is responsible for regulating cell contraction, but it also modulates several currents that affect the action potential. Thus, spontaneous calcium release in the interbeat interval, during diastole, may elicit extra action potential depolarizations and excitation waves, potentially disrupting normal wave propagation. This sometimes leads to the formation of rotors (functional reentry) and eventually a disordered electrical state characteristic of fibrillation [7–9].

Often, this focal activity is due to the presence of periodic calcium waves, that result in calcium oscillations [10–15]. In paced cardiac cells, oscillations necessarily compete with the external pacing frequency and they may be behind occurrences of spontaneous calcium release events during diastole [16]. Calcium oscillations arise typically due to a malfunction of the Ryanodine Receptor (RyR) [16–19], a ligand-gated channel [20] that controls the amplitude of the intracellular calcium transient, by regulating the release of calcium stored at the sarcoplasmic reticulum (SR). Since calcium dynamics in cardiac cells is regulated by the release of calcium at several tens of thousands of RyR clusters (termed calcium release units, CaRUs), global oscillations must appear as a result of an oscillatory regime at the local cluster level that can later be coordinated by diffusion of free calcium. Alternatively, when synchronization is not complete, oscillations at the local level can give rise to periodic calcium waves, providing a pro-arrhythmic substrate [21, 22]. Calcium oscillations have been observed to appear in ventricular myocytes under elevated values of cytosolic calcium [23], due to periodic opening and closing of the RyRs. An increase in cytosolic calcium concentration results in a higher frequency of the oscillations until, at larger values, the SR is depleted because the RyR becomes permanently open [23]. A similar transition has also been studied in models under conditions of SR calcium overload [24–26].

In this paper, we use a detailed subcellular calcium model [27] to show the appearance of periodic calcium waves and then analyze this phenomenon using a deterministic model of calcium in a cardiac cell (or in a CaRU). Within this model, we study the existence and stability of different solutions. We show that oscillations typically appear at high global calcium concentration and/or high RyR open probability. Their appearances depend on a delicate balance between the total calcium level in the cell and the level of buffering of calcium available. For instance, at high values of calsequestrin (CSQ), the system presents a transition from a low concentration, excitable state, to a high concentration state. Such a transition has been proposed to be the basis of complex states, such as long-lasting sparks [28]. At low concentrations of CSQ, in between these two stable states, oscillations appear. We study this transition using a minimal model, that includes the concentration of dyadic and SR calcium and the open probability of the RyR and show that it suffices to explain the appearance of oscillations. A further reduction to a minimal two-dimensional model allows us to explain the transition to the oscillatory regime in terms of the nullcline structure of the system.

## Materials and methods

The methods used in this paper have two clear different natures. First, we use a fully detailed subcellular stochastic model of calcium handling to report numerical results showing calcium oscillations. We analyze under which conditions oscillations appear in a controlled scenario where no external pacing is present, and there are no calcium fluxes with the extracellular medium. Later, to gain insight regarding the origin of the oscillations that we observe in the full model, we construct a minimal deterministic model for the local dynamics of calcium at the level of the Calcium Release Unit. The numerical and mathematical analysis of this model allows us to focus on the substrate of the oscillations disregarding the coordination effects of the full model.

### Detailed subcellular calcium model

We model the spatial structure of the cell as in a previous model of a cardiomyocyte presented in Marchena and Echebarria [27], which has been modified to add the effects of calsequestrin. The equations of the model read:

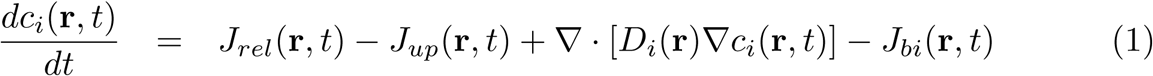

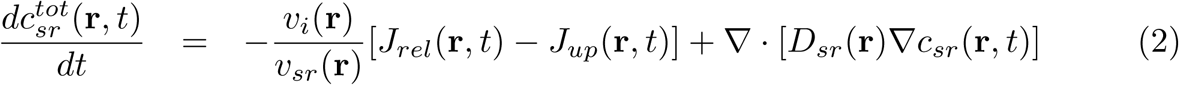

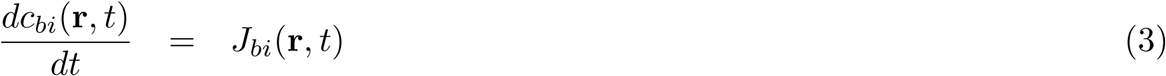

where *c*_*i*_ is the calcium concentration in the cytosol, 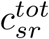 the total calcium concentration in the SR, and *c*_*bi*_ represents the concentration of a given buffer in the cytosol. Besides, *J*_*rel*_ and *J*_*up*_ are the release flux from the SR and the uptake by SERCA, respectively, and *J*_*bi*_ represents the binding of free calcium to the different buffers in the cytosol (TnC, SR binding buffer and CaM). These currents are given by:

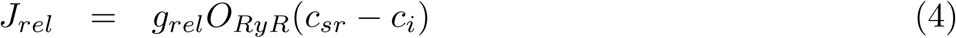

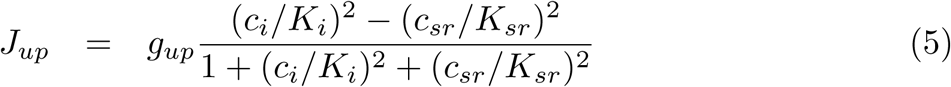

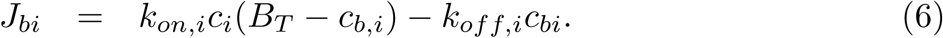

The spatial structure of the model includes cytoplasmic and SR spaces, with a spatial discretization of 100 nm. The volume fraction between cytosolic and SR spaces, *v*_*i*_/*v*_*sr*_, is considered to vary spatially, with different values whether the point is close to the z-line or in the inter z-line space. The release flux *J*_*rel*_ carries Ca^2+^ ions from the SR to the cytoplasm through the RyRs. The RyR channels, indicated by a yellow dot in Fig. 1a, are distributed over the cell along the z-lines with a Gaussian distribution in both transversal and longitudinal axes. A collection of grid points presenting RyRs forms a cluster, i.e., a CaRU. We consider that a CaRU contains 36 RyRs, divided equally among 4 grid points, each one containing 9 RyRs. Each RyR can be in one of four states: open (*O*), close (*C*) and two inactivated states (*I*_1_ and *I*_2_) as it is shown in Fig. 1b. The transitions among these states is considered to be stochastic. In the release flux, the variable *O*_*RyR*_ is the fraction of RyRs that are in the open state and is calculated for all grid points that have a group of RyRs. All the details of the spatial model structure and the values of the parameters can be found in Marchena and Echebarria [27].

**Fig 1.**
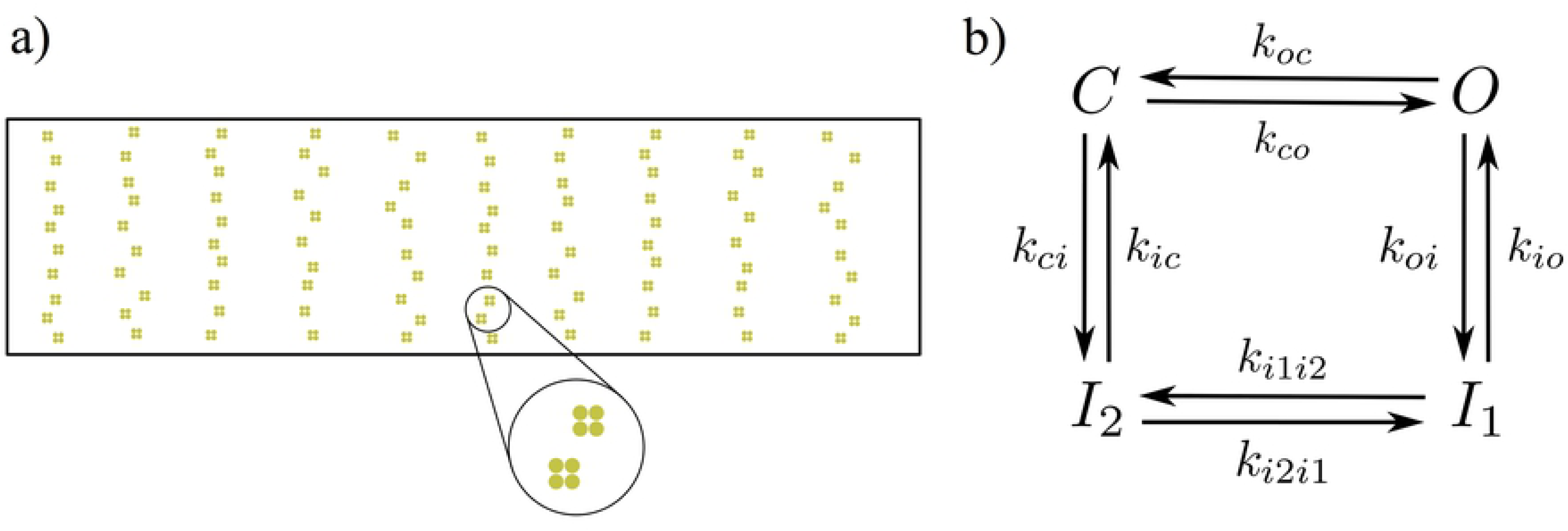
a) RyR distribution in the cell. Each CaRU is formed by four simulation voxels, each one containing 9 RyRs. Thus, all CaRUs are formed by 36 RyRs. The CaRUs are distributed over the cell along the z-lines with a Gaussian distribution in both transversal and longitudinal axes. b) Each RyR follows a four state model, with stochastic transitions among the different states.

Besides the concentration of calcium in the SR and the cytosol, we also consider in the model the concentrations of several buffers. In particular, TnC, CaM and SR-bound buffers in the cytosol [27] and Calsequestrin (CSQ) in the SR. Due to the addition of CSQ in the model, two parameters have been adjusted from the parameters published in Marchena and Echebarria [27]: the opening rate parameter, now *k*_*a*_ = 2.1 · 10^−3^ *µ*M^−2^ms^−1^, and the dependence of the open probability of the RyR on luminal calcium, now *EC*_50−*SR*_ = 450 *µ*M. Contrary to the buffers in the cytosol, the dynamics of CSQ is considered to be fast [29–31] compared with the release time scale. If we denote by *c*_*bSQ*_ the calcium concentration bound to CSQ in the SR, then, the amount of bound calcium is given by:

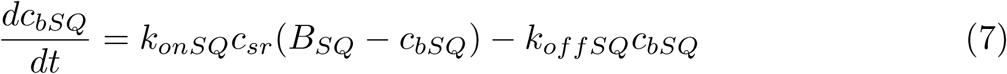

Assuming fast binding, the stationary condition for *c*_*bSQ*_ (*ċ*_*bSQ*_ = 0) is:

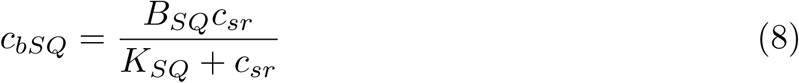

where *K*_*SQ*_ = *k*_*offSQ*_/*k*_*onSQ*_ is the dissociation constant. From this, the concentration of free calcium can be obtained solving Eq. (2) for the total amount of calcium in the SR, 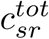

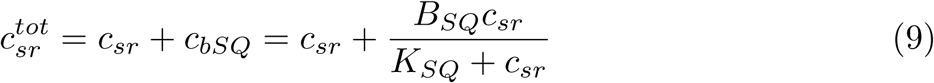

Solving this equation we obtain the value of free calcium in the SR,

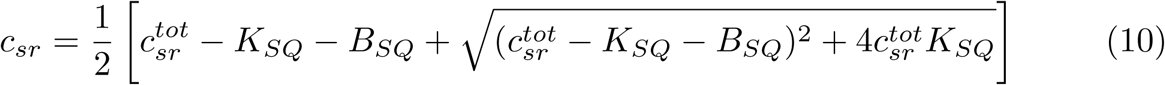

The advantage of using this formulation of the rapid buffer approximation over the more usual, for example in [32], is that it conserves mass exactly.

Under physiological conditions, the total amount of calcium in the cell at steady state is fixed by calcium homeostasis, i.e. the complex interaction of LCC, exchanger, and pumps, which affect the steady state level at which the calcium entering the cell balances the calcium extruding. In this work, we are interested in studying instabilities in calcium cycling, under constant cell calcium content. This allows us to focus the analysis on the conditions for the appearance of calcium oscillations under different possible calcium homeostatic levels. Thus, we neglect calcium exchange with the extracellular medium, setting the conductances of the L-type calcium channels and the NCX equal to zero. Then, the total amount of calcium in the cell, *Q*_*T*_, is given by:

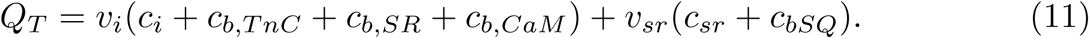

For a better comparison with the results from a reduced calcium model, described later, we will consider as our control parameter the average calcium content of the cell 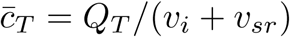. Thus, in our simulations, 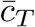 is a constant value that is determined by the initial conditions for cytosolic and luminal calcium (free and bound to buffers).

### Reduced calcium model

The minimal model for the local dynamics of calcium is based on the schematics shown in Fig. 2. We consider a simplified description of the system, with dynamics of the total calcium concentration in the SR, 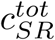, and in the cytosolic space close to the RyR2, or dyadic space, *c*_*d*_, and of the open probability of the RyR, *P*_*o*_,

**Fig 2.**
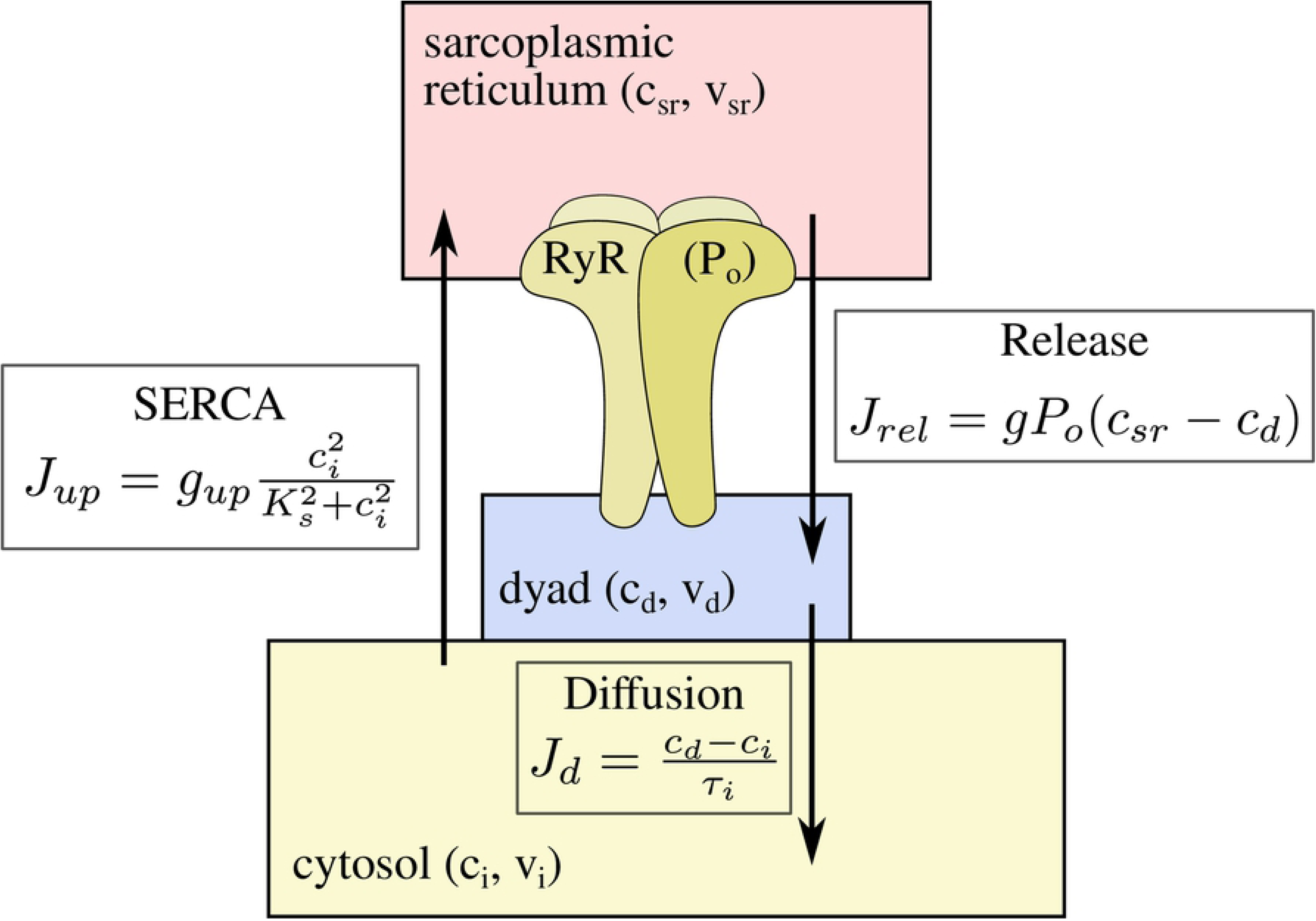
Sketch of the different compartments considered in the simplified model, with the internal variables and the equations of the respective calcium fluxes.

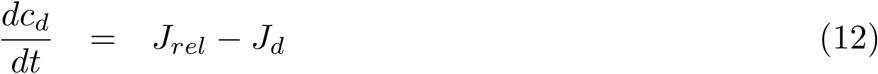

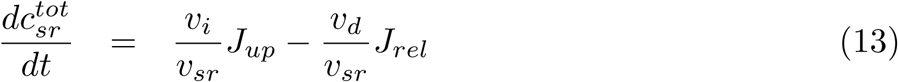

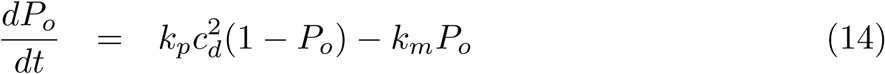

with the currents given by

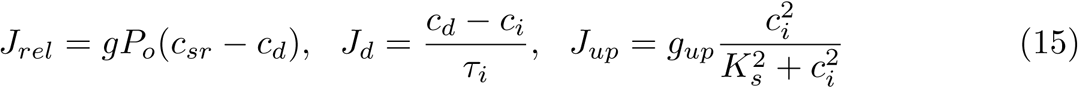

A detailed derivation of these equations and their range of validity can be found in Appendix A. Notice that, as in the full model, we consider a situation where no external pacing is imposed. In this sense, neither external intake from the LCC is considered, nor any extrusion via the sodium-calcium exchanger.

For simplicity, we consider a SERCA pump without an equilibrium condition, that always pumps calcium from the cytosol to the SR. This gives a basal solution at *c*_*i*_ = *c*_*d*_ = 0, instead of the physiological value of ∼ 100nM. However, given that, at basal conditions, *c*_*sr*_ ∼ 1mM, this is a reasonable simplification. As in the detailed subcellular model, we assume the approximation of rapid CSQ buffer, so we can compute the amount of free luminal calcium *c*_*sr*_ from the total luminal calcium 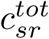 from Eq. (9).

To close the system we should introduce an extra equation for calcium concentration in the cytosol *c*_*i*_. However, as we assume that the total calcium content in the cell is constant, then we have a conservation equation. Therefore, we can compute *c*_*i*_ solving the following quadratic equation for the conservation of 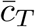

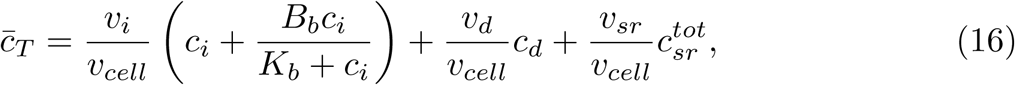

where *v*_*cell*_ is the unit volume defined as *v*_*cell*_ ≡ *v*_*i*_ + *v*_*sr*_ + *v*_*d*_, and *B*_*b*_ is the concentration of a generic buffer in the cytosol.

To simplify the analysis, we proceed to work with the assumption that the dynamics of the RyRs is faster than that of calcium concentration 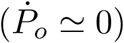, obtaining then a minimal two-variable model. This will be our base-line minimal model. However, we will later also consider an alternative model with fast dynamics for the dyadic calcium concentration (*ċ*_*d*_ ≃ 0). A third possibility, with fast dynamics of luminal calcium, although theoretically possible, does not have much physiological sense, as SERCA is typically slow compared to release or diffusion from the dyadic space.

#### Fast RyR dynamics

In this case, we assume that the open and close dynamics of the RyR are fast, so we can assume that it is in a quasi-steady state 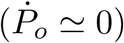. Then, from Eq. (14), we obtain:

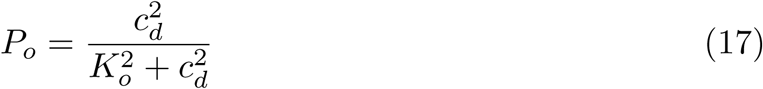

where the parameter 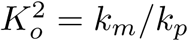 is the ratio of the open and close rates of the RyR.

Then, with these assumptions, the simplified model becomes

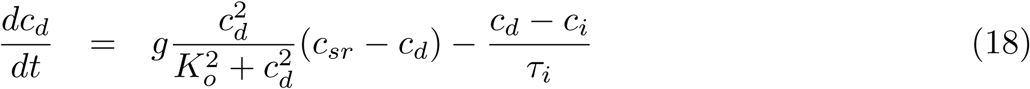

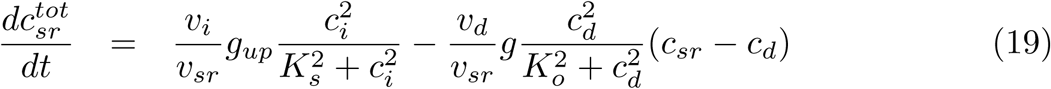

#### Fast dyadic calcium dynamics

In order to test the robustness of the analysis, we also consider a simplified model given by Eqs. (12)-(14), in the limit of fast dynamics in the dyadic space and take *ċ*_*d*_ ≃ 0. Then, from Eq. (12):

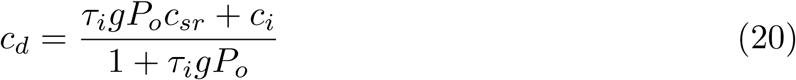

Substituting this expression in Eqs. (13) and (14), we obtain another minimal model, given by

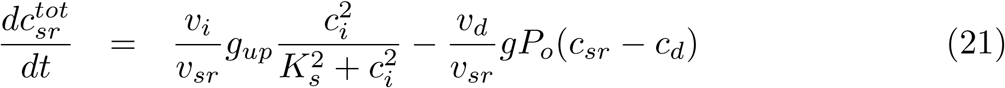

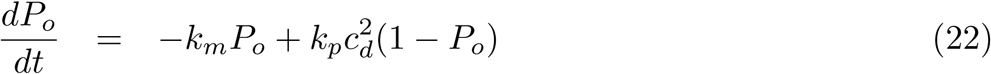

where again, *c*_*i*_ must be computed solving the quadratic equation for the conservation of mass 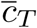 [Eq. (16)]. For simplicity we will consider the case when no calsequestrin is present *B*_*SQ*_ = 0.

## Results

We first present the results of the numerical simulations of both the full detailed model and the minimal model of calcium cycling. Both produce the same basic scenarios for intracellular calcium dynamics, with three different dynamical behaviors, which we then proceed to analyze. The goal of the development of the minimal model is, precisely, to be able to perform this analytical treatment and check how the behavior depends on total calcium and buffering levels.

### Subcellular model

The full detailed model allows us to investigate the different behaviors present in cardiomyocyte calcium cycling when there is no external pacing. We should point out that, under these conditions, the average calcium concentration in the cell 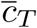 is conserved since the total amount of calcium *Q*_*T*_ in the cell is constant. We produce simulations with different levels of average calcium concentration and observe very different behaviors (Fig. 3a). For the lowest value of 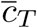, the RyR remains almost closed, and most of the calcium content is stored in the SR. Despite the stochasticity of the system, the average values obtained are reproduced reliably with only the presence of local sparks as fluctuations of this global state. This state corresponds to an excitable state, which is the expected behavior of the cell if it has to react properly to external excitation. We call this general state a global shutdown state.

**Fig 3.**
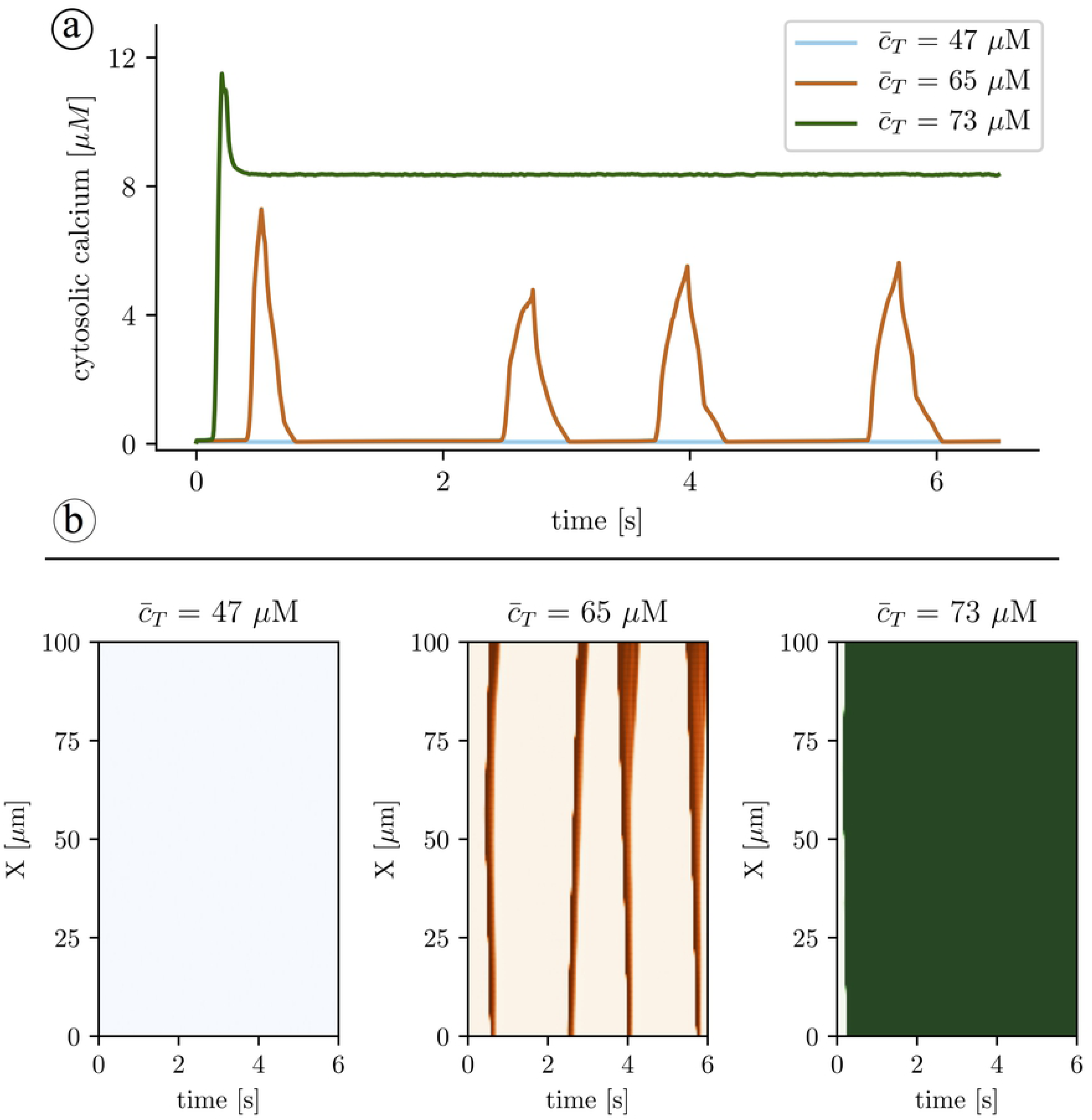
A: Calcium traces obtained with the full subcellular model and three different values of the average calcium concentration, 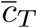. B: Line-scans at different values of the load. Increasing the load, the system undergoes a transition from a low cytosolic calcium state (at 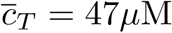), where RyRs remain in the closed state, to spontaneous oscillations, giving rise to calcium waves 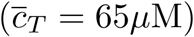. Finally, at high calcium loads 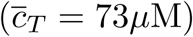 oscillations give rise to a high cytosolic calcium state, where the RyRs remain open, resulting in SR calcium depletion.

When the calcium load increases, the system starts to spontaneously show calcium waves. These waves persist in time with different shapes and durations, giving rise to a nearly periodic oscillation in the global calcium signal. Roughly, we observe one calcium wave per second (Fig. 4). Waves are normally initiated at different sites each time but they appear systematically indicating a strong oscillation at the substrate level that we will address in the discussion.

**Fig 4.**
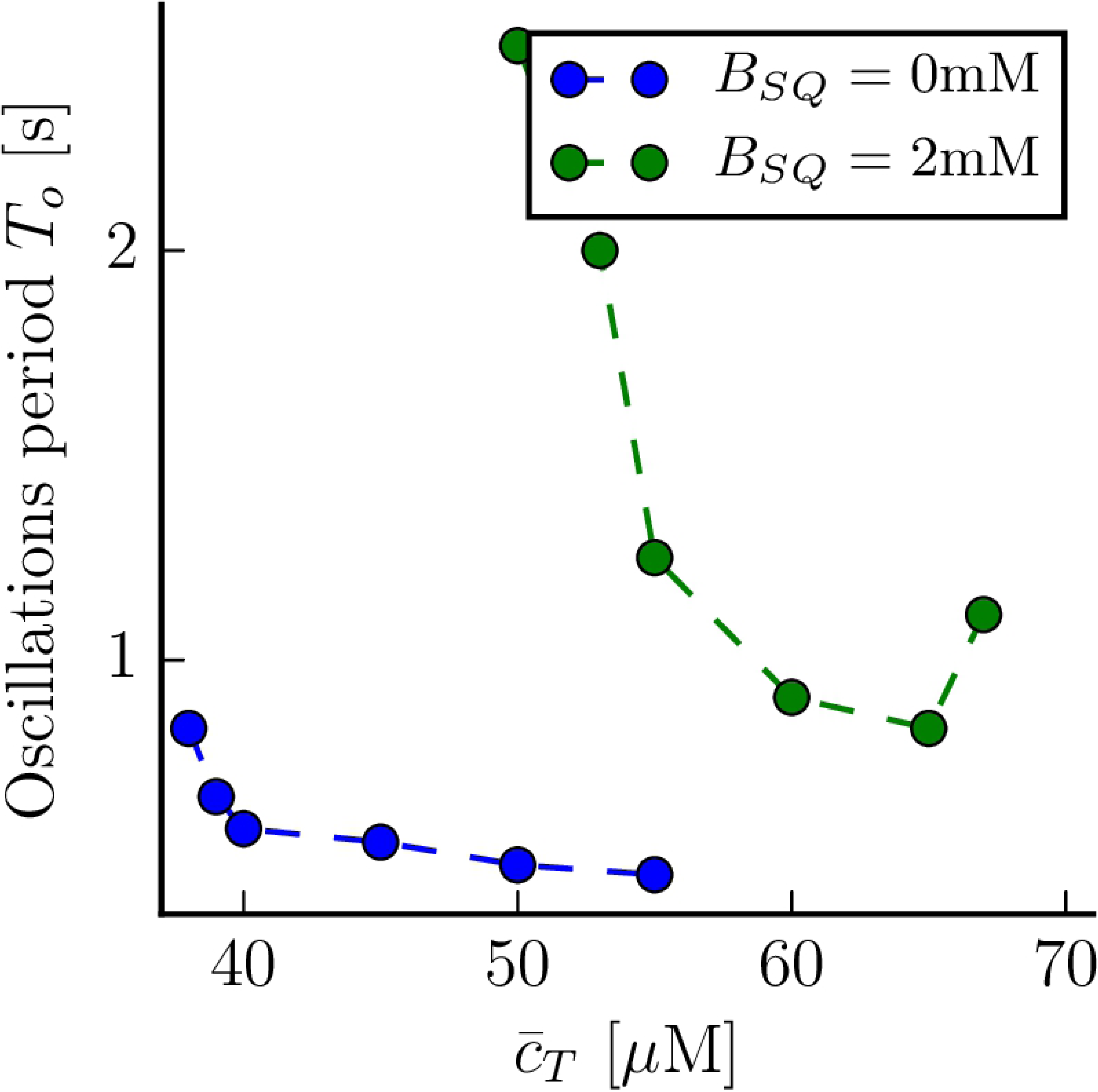
The average period of oscillations at different values of the average calcium concentration 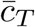, for a concentration of CSQ of *B*_*SQ*_ = 2mM (green dots), and in the absence of CSQ (blue dots).

Finally, at large values of 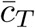 oscillations disappear, giving place to a stable state with a high level of calcium in the cytosolic space and a depleted SR. In this state, the RyRs are generally open allowing for the depletion of the SR and the increase of cytosolic calcium. Except for local fluctuations this state is globally stable and we can call it the open state. This state would not respond to external pacing, however, it would produce the activation of the NCX exchanger which would slowly decrease the average concentration model. As we pointed out previously, the elimination of the calcium intake and extrusion in the model allows us to focus on the general behavior of the cell under different homeostatic scenarios. Numerical simulations indicate that as the calcium level is increased, the cell goes from a shut-down and ready-to-respond state to an oscillatory regime to a global open state where the cell does not respond.

We must point out that a similar trend has been observed experimentally by Stevens et al [23], even if in the experimental preparation the control parameter was the amount of cytosolic calcium, and not total calcium, as in our simulations. Oscillations appear as the amount of calcium in the cell increases, giving rise to a state with depleted SR calcium (and RyRs in the open state), at high calcium concentrations. Furthermore, experimentally it has been shown that changes in buffering levels can have also important effects on this transition. More specifically, Stevens et. al [23] have shown that the reduction of CSQ in the SR bulk enhances the appearance of oscillations. We have checked if this situation is also present in our simulation and found this to be the case. As shown in Fig. 4, when we reduce the CSQ concentration, the oscillations appear at lower values of 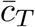, they have a higher frequency and the range of oscillations in terms of 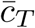 becomes broader.

### Minimal model without calsequestrin

We proceed to explain the results obtained in the minimal model where we find the same qualitative behavior as in the results obtained with the full subcellular stochastic model. We have performed simulations in the approximation of fast RyR dynamics at different values of the cell average calcium concentration 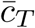. We consider first the case when no calsequestrin is present, *B*_*SQ*_ = 0. As we observed in the full subcellular model, at low values of 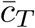 the system remains in a low concentration steady state (Fig. 5). In this state the system is excitable, so the fixed point is locally stable, but a large enough perturbation produces an increase in calcium concentration, resulting in a calcium transient typical from CICR. On the other hand, at high calcium loads, there is a new fixed point with high cytosolic calcium concentration, that has been related to the appearance of long-lasting sparks [28]. As in the subcellular model, in between the low concentration fixed point and the permanently open state, the system presents oscillations (Fig. 5), that are stable for a quite broad range of loads.

**Fig 5.**
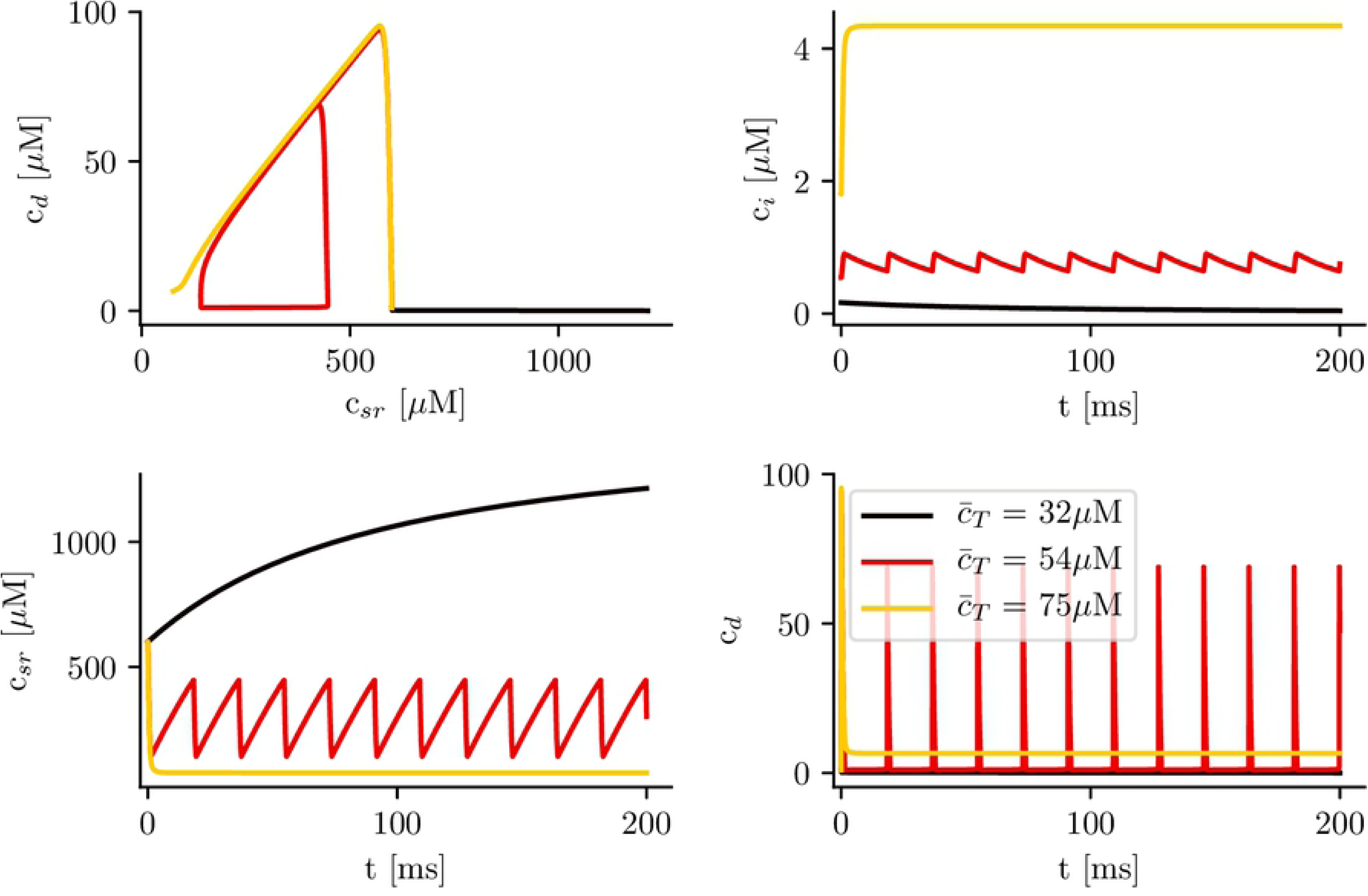
Time traces of the different calcium concentrations for different values of total calcium concentration, 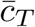, and calsequestrin concentration set to zero (*B*_*SQ*_ = 0). After a transient, the system ends up in either a steady state which is excitatory at low levels of total calcium in the cell with observed low levels of calcium in the cytosol, in an oscillatory state with intermediate levels of total calcium in the cell, or in a state of high total levels of calcium in the cell with observed high cytosolic calcium levels.

Indeed, the number of stationary solutions changes with the calcium concentration 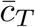. The fixed points of the system can be found from:

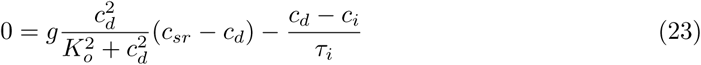

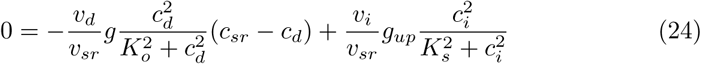

together with:

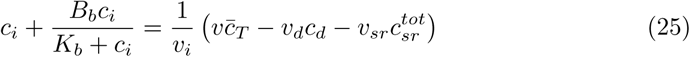

Eqs. (23)-(25) represent three algebraic equations that give the concentrations *c*_*i*_, *c*_*d*_ and *c*_*sr*_ as a function of total calcium concentration in the cell 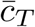. When no calsequestrin is present, *B*_*SQ*_ = 0, it is easy to obtain that

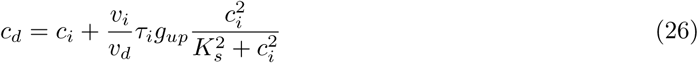

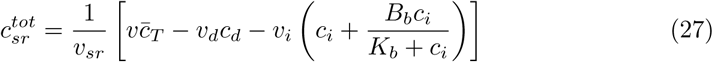

Introducing these expressions into Eq. (23) we obtain an equation of the form 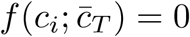. For each value of 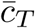 we can obtain the values of *c*_*i*_ that solve the equation. For instance, for a global average calcium concentration of 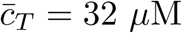 we have only one solution, as shown in Fig. 6a. This solution, given by *c*_*d*_ = *c*_*i*_ = 0 and 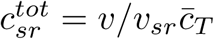, exists for all values of 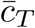. At high values of the average concentration, 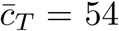 and 75 *µ*M, another two solutions appear, as depicted in Figs. 6b and c.

**Fig 6.**
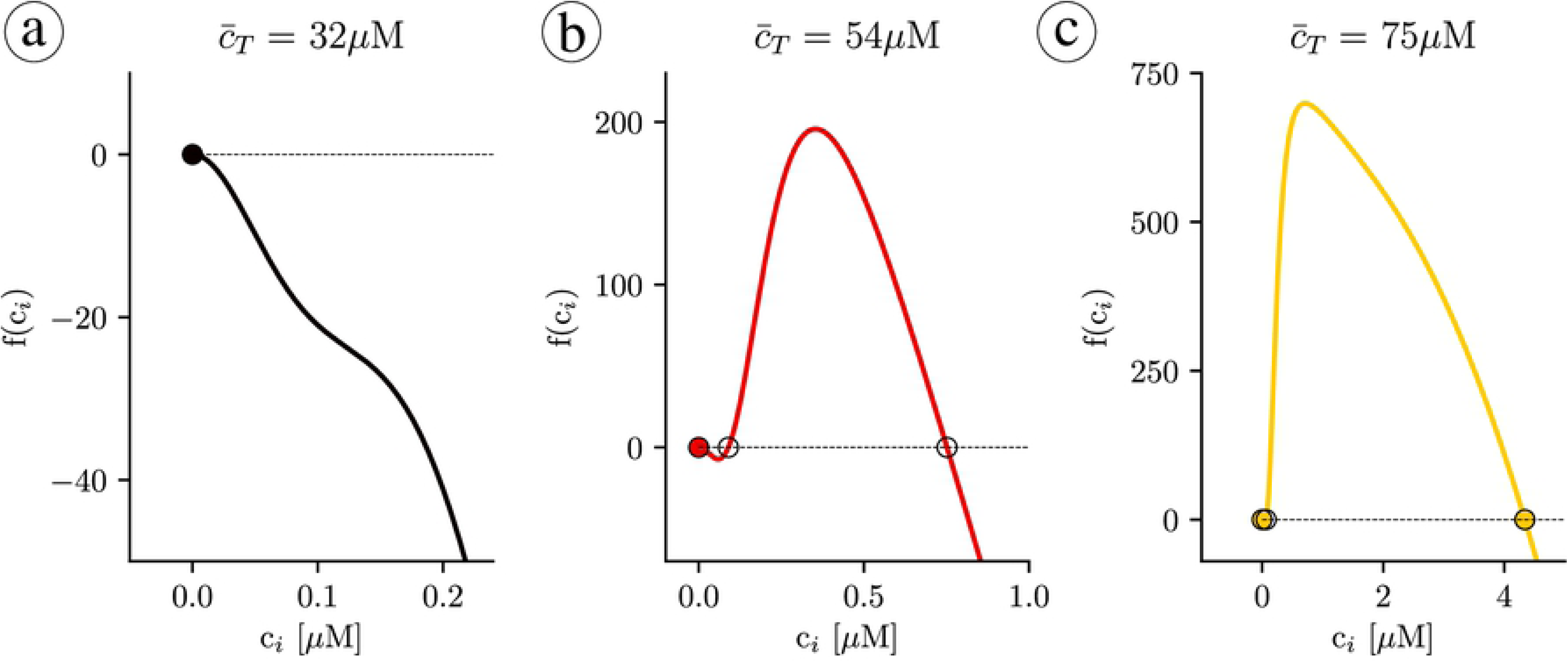
Plot of the function *f* (*c*_*i*_) for different values of the 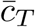 concentration. a) At low concentrations there is a single fixed point. b) At higher concentrations two extra unstable fixed points appear. c) At high concentrations the upper fixed point becomes stable.

To calculate the stability of the stationary solutions, we have computed the value of the eigenvalues of the Jacobian matrix, corresponding to Eqs. (18)-(19). We find that, while the lower branch is always stable, the other branch of solutions is unstable for a large range of parameters (Fig. 7), due to the appearance of oscillations. The stability of the corresponding periodic orbit has been calculated using XPPAUT [33] (Fig. 8). We obtain that, as 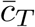 is increased, a limit cycle appears in a global homoclinic bifurcation, with zero frequency (Fig. 8c). Increasing 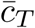, this limit cycle finally disappears in a Hopf bifurcation, at which the upper fixed point becomes stable (Fig. 8b).

**Fig 7.**
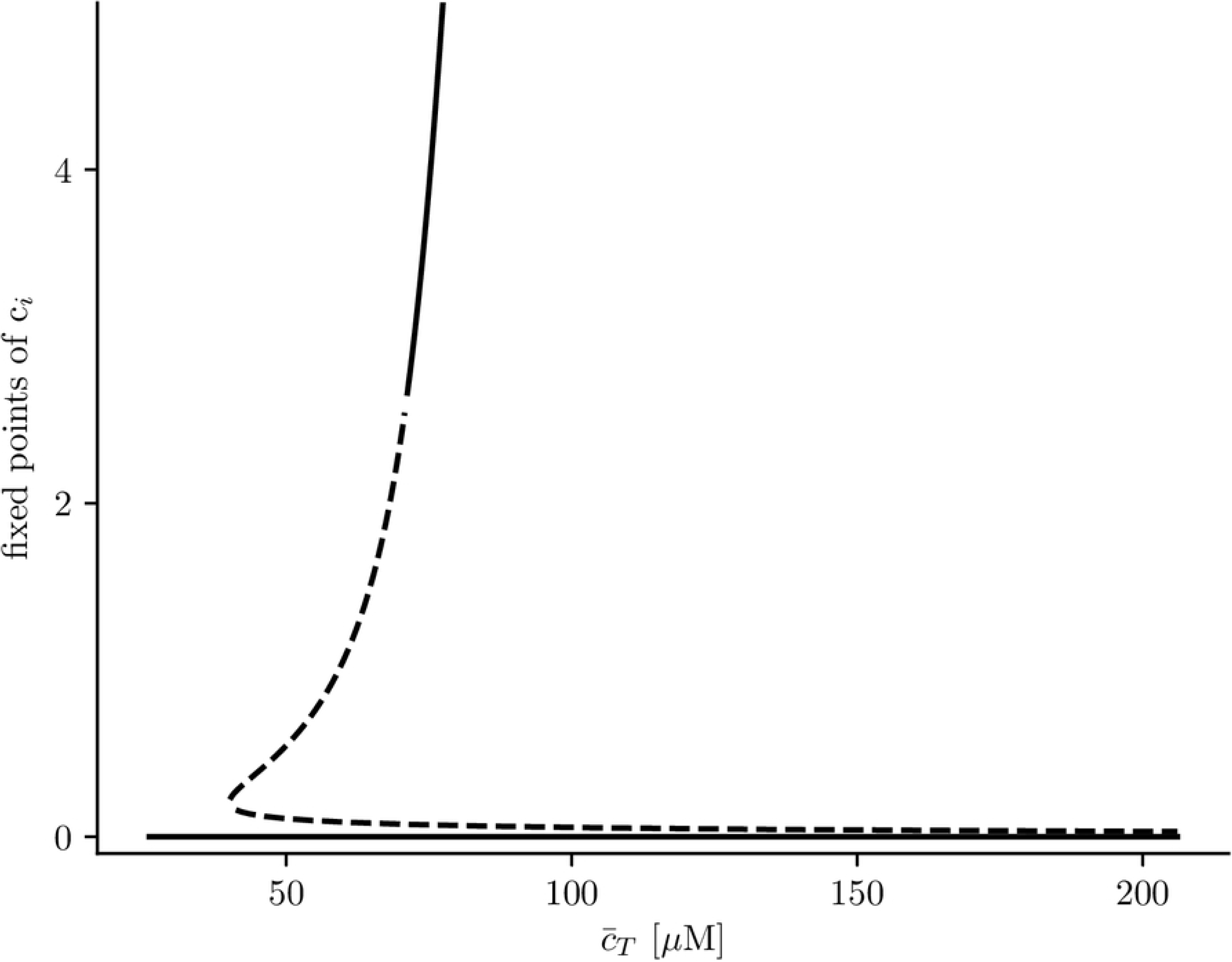
Solutions for cytosolic calcium concentration, *c*_*i*_, as a function of total calcium concentration, 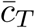. Discontinuous lines represent unstable solutions while continuous lines stable ones.

**Fig 8.**
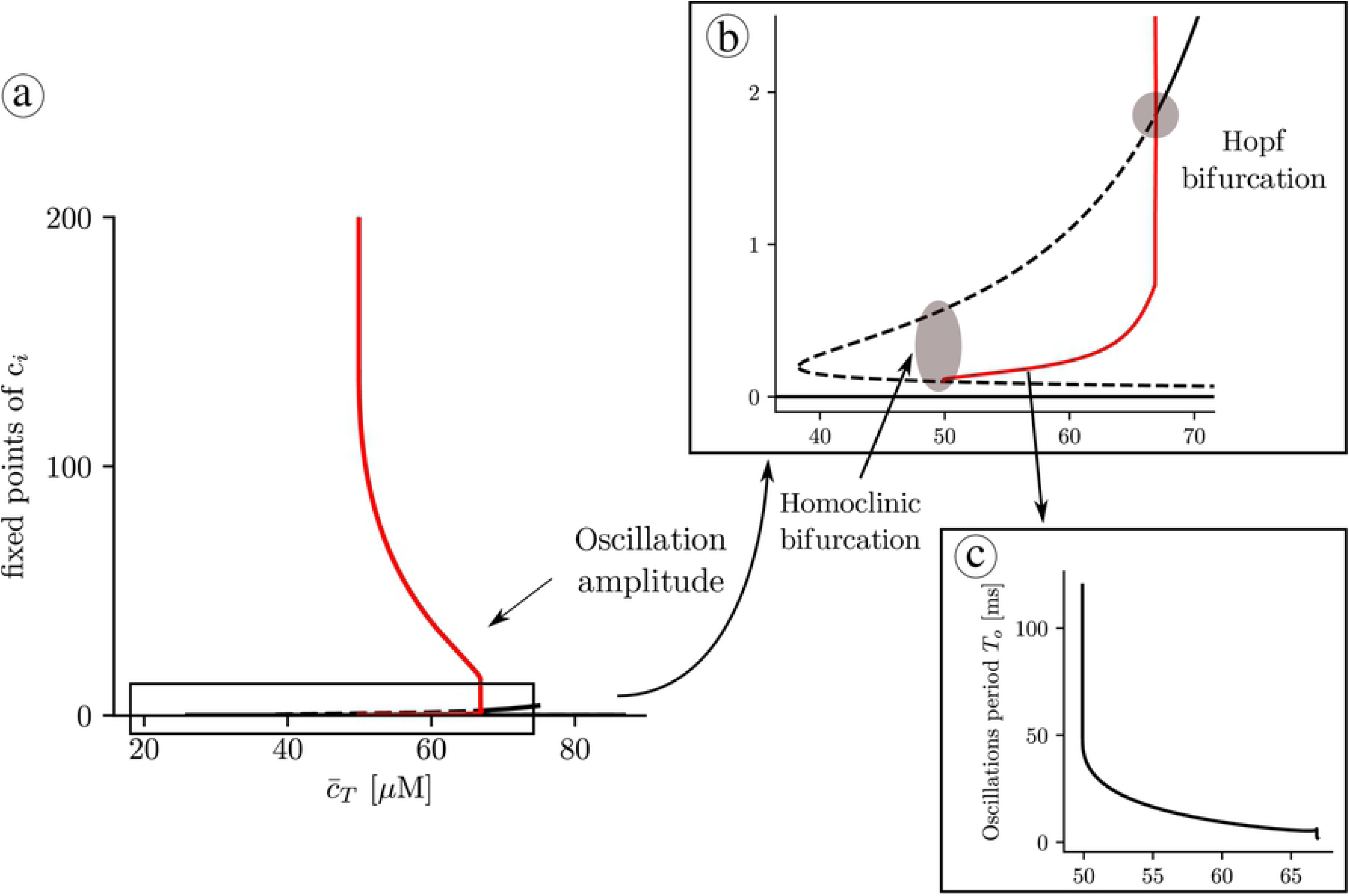
a) Solutions for cytosolic calcium concentration, *c*_*i*_, as a function of total calcium concentration, 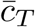. Discontinuous lines represent unstable solutions and continuous lines stable ones. When reducing the total concentration, at 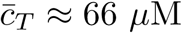, a limit cycle emerges in a Hopf bifurcation, from the upper state, that then becomes unstable. The red lines represent the lower and upper values of the limit cycle. At, 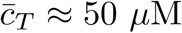, the intermediate unstable fixed point collides with the limit cycle, that disappears in a homoclinic bifurcation. Below 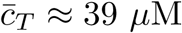, the RyR close state is the only solution. c) Oscillation periods as a function of 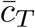.

### Minimal model with calsequestrin

We present now the main results of the simulation in the fast RyR minimal model when calsequestrin is present. We use Eqs. (18)-(19), with Eq. (9) that relates free with CSQ-bound SR calcium concentrations. Typically *K*_*SQ*_ = 650*µM* and *B*_*SQ*_ somewhere between zero and 20 mM. We have then calculated the different fixed points as a function of the total concentration 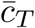 for different values of *B*_*SQ*_ (Fig. 9). When *B*_*SQ*_ *≠* 0, increasing 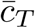, the appearance of two extra solutions occurs at larger values of 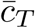, meaning that the close state solution is stable for a wider range of 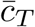. Besides, the oscillatory range becomes narrower when *B*_*SQ*_ increases, until at certain point oscillations disappear. The disappearance of oscillations is due to the transformation of the bifurcation at which the upper fixed value gains stability, from a Hopf bifurcation into a saddle-node bifurcation.

**Fig 9.**
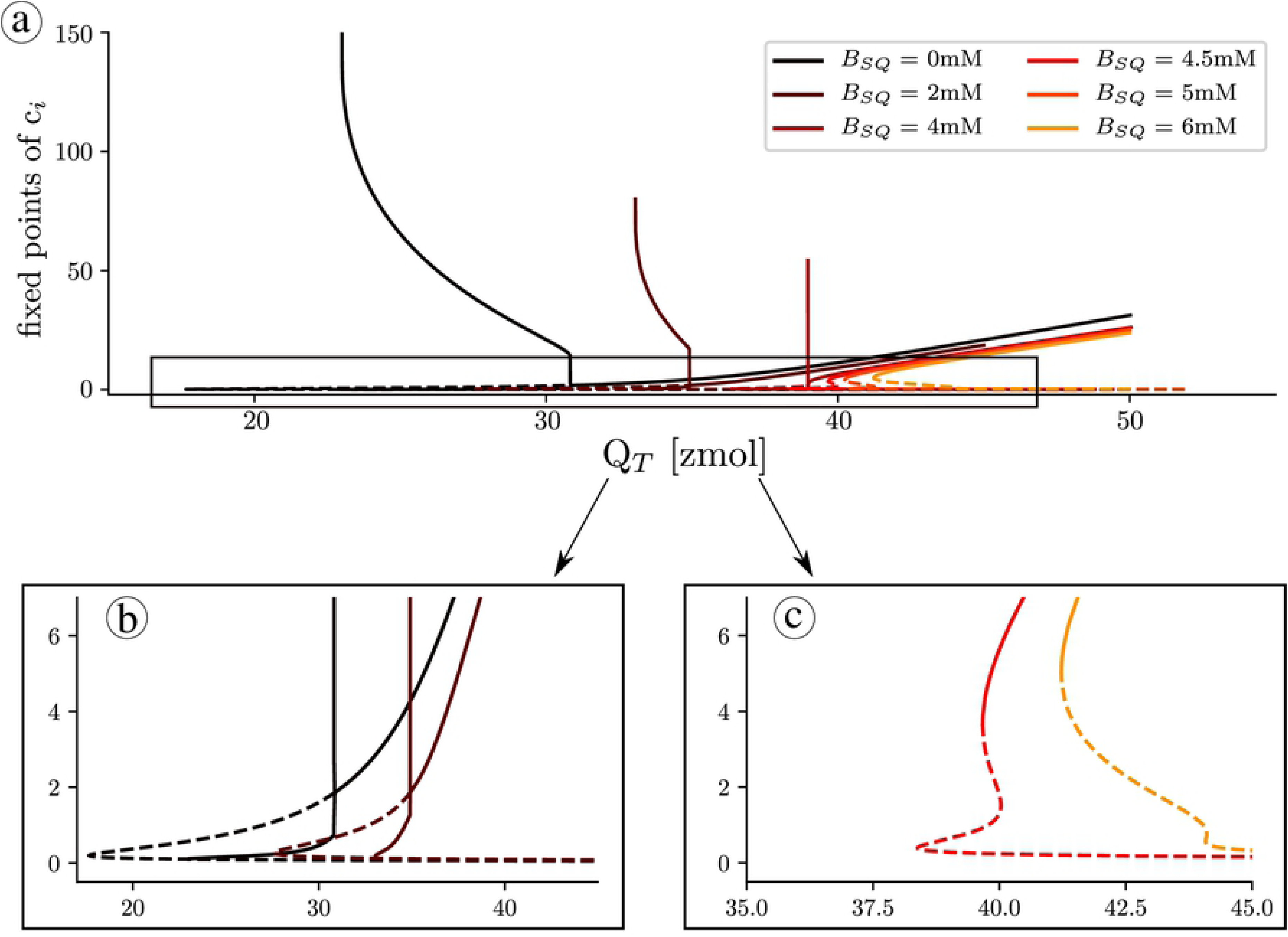
Number of fixed points and stability of those fixed points as a function of total calcium in the unit, for different values of calsequestrin concentration.

In addition, at this point, the system presents five fixed points where only the lowest and uppermost are stable. It is important to note that this shows that CSQ enhances the elimination of oscillations. Finally, for high values of 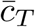, the system has again three fixed points.

### Analytical results

The mathematical tractability of the minimal model allows us to get a better understanding of the transition to the upper state via oscillations. A first insight can be obtained by plotting the nullclines of the system (Fig. 10), which can help us understand the main mechanisms behind the transition to the oscillatory state. In particular, we obtain the critical average calcium concentration for the onset of oscillations, which depends on buffering levels, and the conditions for the appearance of the upper state.

**Fig 10.**
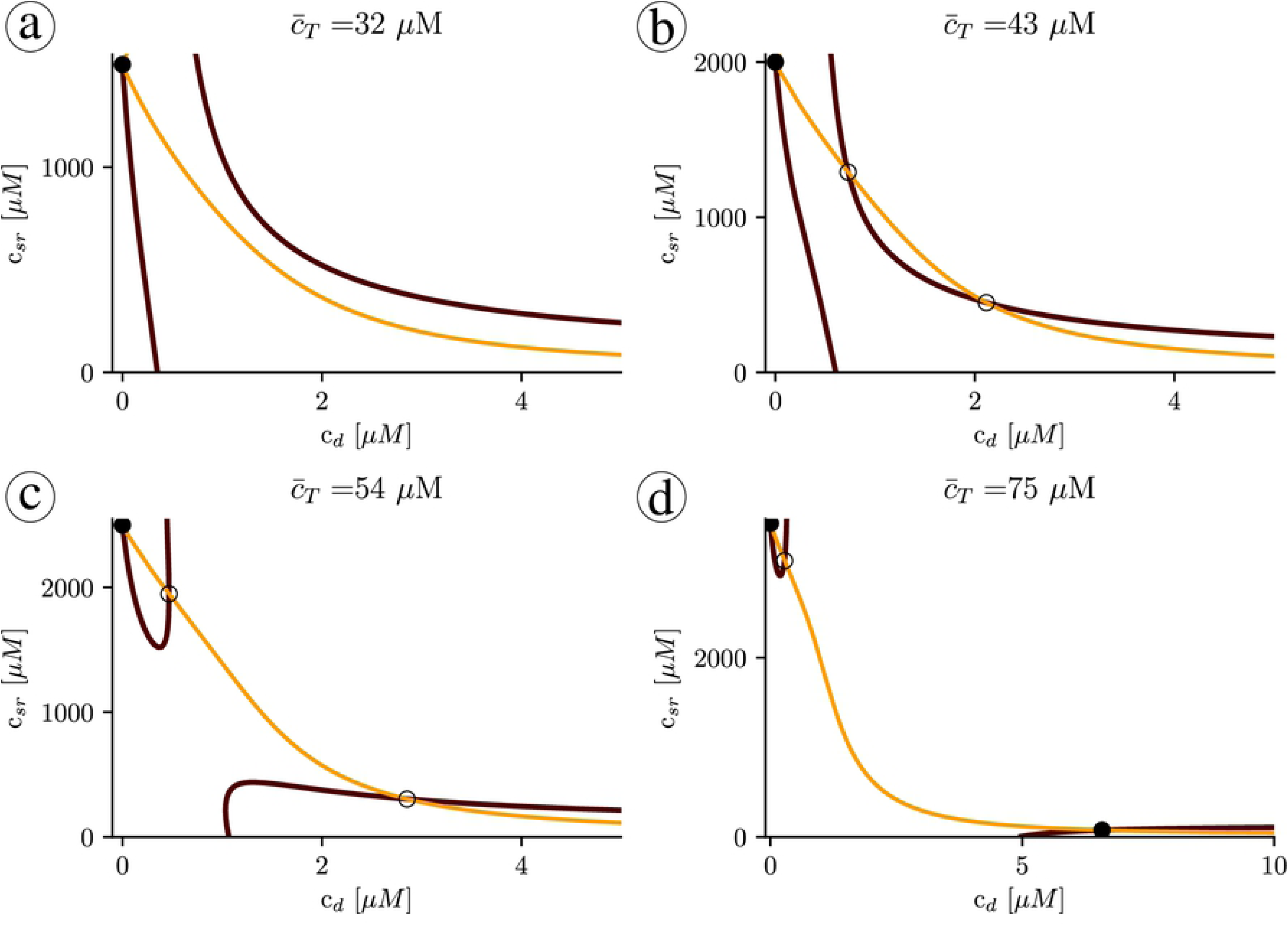
Structure of the nullclines at different values of 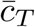 indicated in the title of each panel. The black line indicates the first nullcline *ċ*_*d*_ = 0, while the orange lines corresponds to *ċ*_*sr*_ = 0. Dots indicate the fixed points. Filled dot: stable fixed point and unfilled dot: unstable fixed point.

#### Nullclines and stability of solutions

Besides the transition from one to three solutions (Figs. 10a and b), nullclines present a clear restructuring of their branches well before the upper state becomes stable. Increasing the total concentration, there is a sudden pinch-off in the *c*_*d*_-nullcline (Figs. 10b and c). Before this change in nullcline topology, the lower state (with all the calcium in the luminal space and none in the cytosol) is the only stable attractor. Once the *c*_*d*_-nullclines split, oscillations may appear around the upper unstable fixed point.

We can understand the effect of the pinch-off, plotting the trajectories for values of 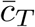 close to the transition. Below the pinch-off, the trajectory follows the fast dynamics of the *c*_*d*_-nullcline (the black line in Figs. 10 and 11), until it reaches the fixed point. As the load is increased, the branches of the *c*_*d*_-nullclines come closer until at a certain point the break-up occurs (Fig. 11c). Due to the emergence of the pinch-off, the system dynamics follows the lower nullcline up to the tip, at the largest value of *c*_*sr*_, where, due to the fast dynamics in the *c*_*d*_ direction, it jumps to follow again the nullcline, at larger values of *c*_*i*_. Since there is no stable point, the trajectory starts a persistent oscillation around the unstable fixed point. We can state that, for this problem, the nullcline break-up is the necessary and sufficient condition to obtain oscillations.

**Fig 11.**
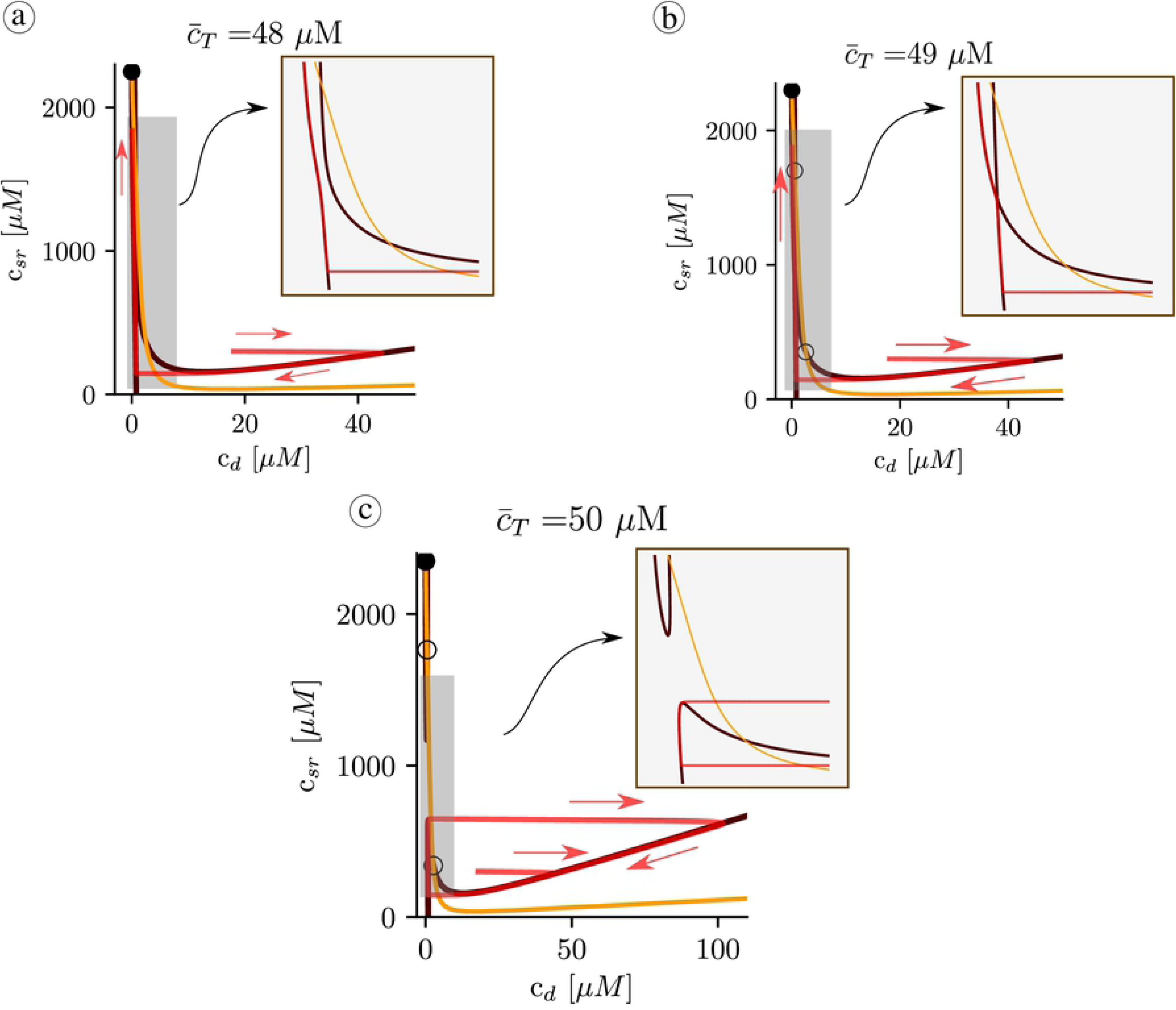
Structure of the nullclines at different values of 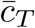 indicated in the title of each panel. The black line indicates the nullcline *ċ*_*d*_ = 0, while the orange line corresponds to *ċ*_*sr*_ = 0. The red curve is a trajectory with a direction indicated by the red arrows.

#### Onset of oscillations

Using this observation, we can calculate analytically the critical value of 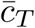 beyond which the system oscillates. At the pinch-off of the nullclines, the cubic solution of *ċ*_*d*_ = 0 (black nullcline) loses two of the three solutions at a given *c*_*sr*_. To calculate this point analytically, first we observe that the pinch-off occurs at values of *c*_*d*_ *< K*_*o*_ (*K*_*o*_ = 15*µ*M). To simplify the calculations, let us make the approximations that, at the pinch off, the *c*_*d*_ satisfies *c*_*d*_ ≪*K*_*o*_ and *c*_*d*_ ≪*c*_*sr*_. Being this the case, then Eq. (18) reduces to

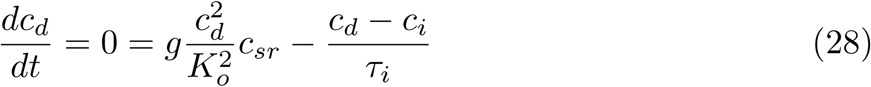

Furthermore, from Eq. (25), we can write *c*_*i*_ in terms of *c*_*sr*_ (assuming *B*_*SQ*_ = 0, *c*_*d*_ ≪ *c*_*sr*_)

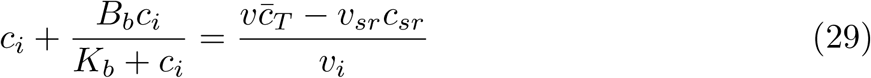

Assuming that the concentration of bound cytosolic calcium is much larger than that of free cytosolic calcium, we obtain

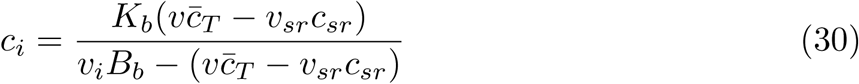

From the polynomial equation for *c*_*d*_, Eq. (28), solutions of *c*_*d*_ are lost at values of *c*_*sr*_ given by 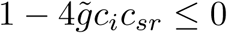, with 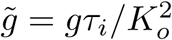. Expanding, this gives the critical value of 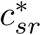 as:

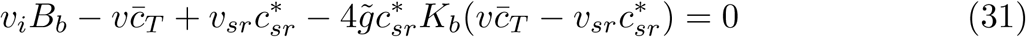

The same equation can be written as

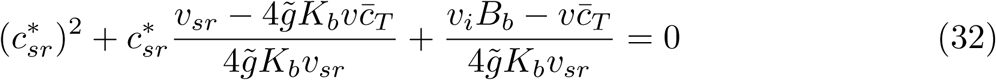

Once the pinch-off has been produced (Fig. 10c), there are two values of 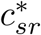, corresponding to the lowest and highest values of the upper and lower nullclines, respectively. Just at the pinch-off these two points merge. This allows to establish the critical value at which the oscillation appears as the one that makes zero the discriminant of the second order polynomial of 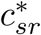. After some algebra, the condition for the critical 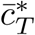 becomes

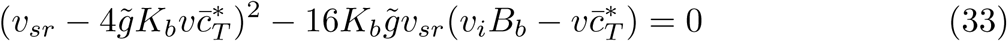

which can be written as

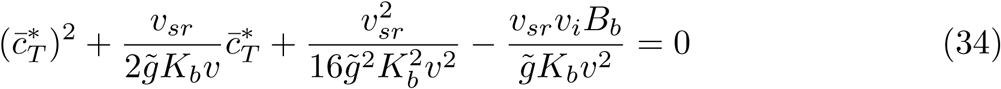

This gives

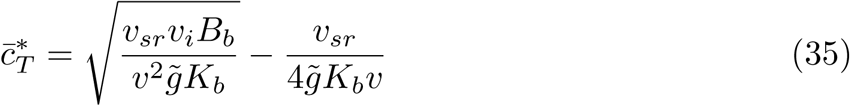

Using the parameters in Table 1, this expression gives a critical value for the onset of oscillations at around 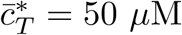, that given the approximations considered, agrees quite well with the value obtain from the simulations of 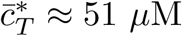. Thus, when the total calcium content exceeds this critical value 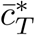, corresponding to a calcium concentration in the SR of 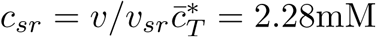 (in the lower state), the system starts to oscillate, at a value of diastolic SR calcium load, given by:

**Table 1.** Parameters of the model.

| Parameters |  |  |
| --- | --- | --- |
| Parameter | Units | Value |
| $v_d$ | $\mu m^3$ | 0.001 |
| $v_i$ | $\mu m^3$ | 0.45 |
| $v_{sr}$ | $\mu m^3$ | 0.01 |
| $\tau_i$ | $ms$ | 0.01 |
| $g_{up}$ | $\mu M/ms$ | 0.5 |
| $K_s$ | $\mu M$ | 0.2 |
| $g$ | $ms^{-1}$ | 20 |
| $k_p$ | $\mu M^{-2}ms^{-1}$ | $10^{-3}$ |
| $k_m$ | $ms^{-1}$ | 0.25 |
| $K_o$ | $\mu M$ | 15 |
| $B_b$ | $\mu M$ | 80 |
| $K_b$ | $\mu M$ | 0.5 |

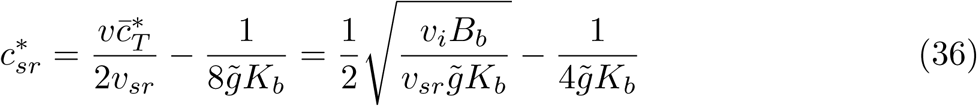

which gives a value of 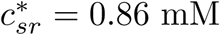 mM. At the onset of oscillations, there is thus a sudden decrease in basal SR calcium concentration, to less than half its previous value before the oscillations.

An increase in the quantity of cytosolic buffers (higher *B*_*b*_) results in a delay in the onset of oscillations, that would occur at higher calcium load (Fig. 12a). A higher calcium affinity (lower *K*_*b*_) on the contrary, would result in oscillations at lower loads (Fig. 12b). It is interesting to notice, too, that the strength of SERCA does not affect the onset of oscillations.

**Fig 12.**
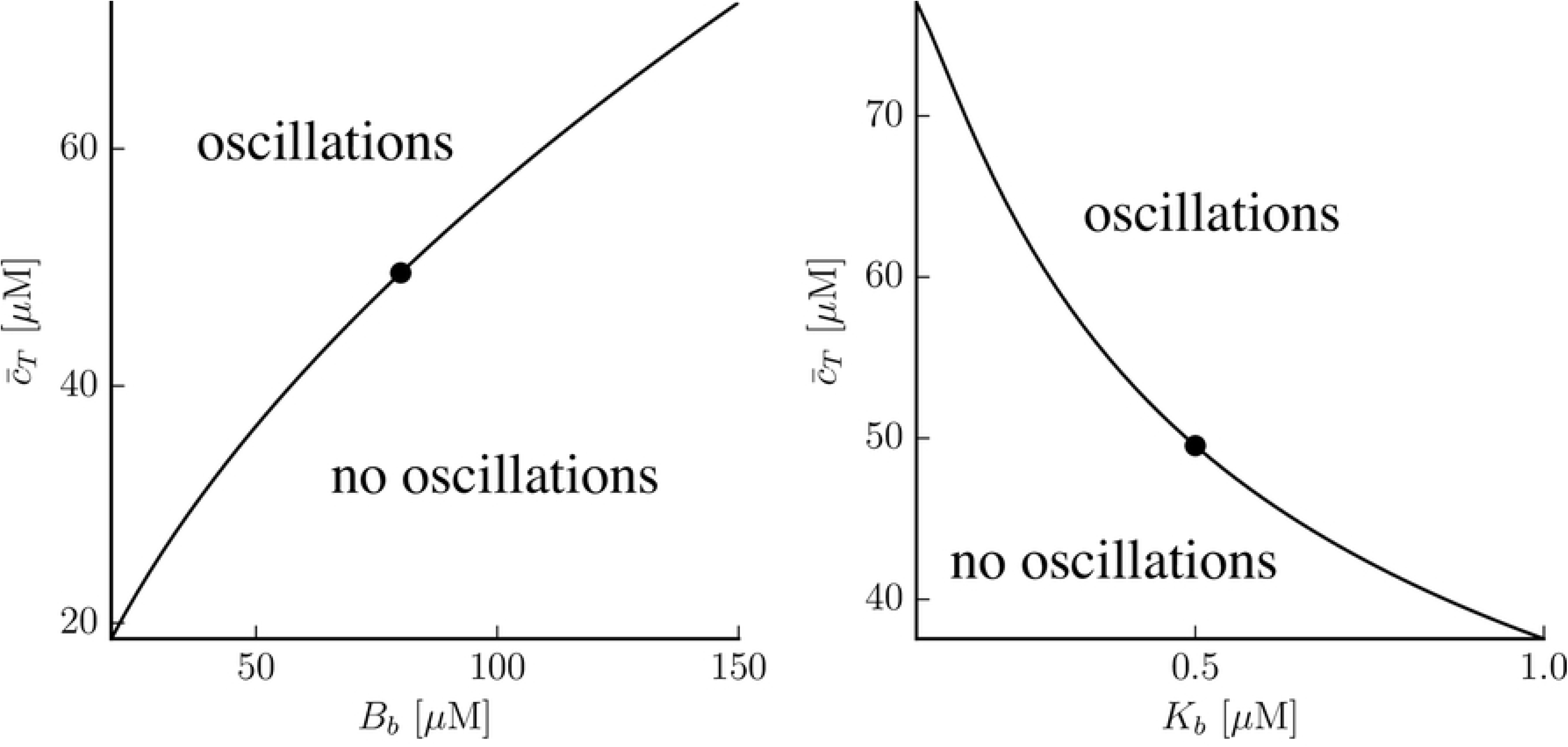
Dependency of the onset of oscillations with buffer parameters. The filled dots represent the control values *B*_*b*_ = 80*µ*M, *K*_*b*_ = 0.5*µ*M, given in Table 1.

#### Transition to the upper state

The oscillations disappear at a Hopf bifurcation when the upper state becomes stable. It is possible to relate this transition to the structure of the nullclines in Fig. 10. For that, let us recover the definition of the Jacobian matrix **J**:

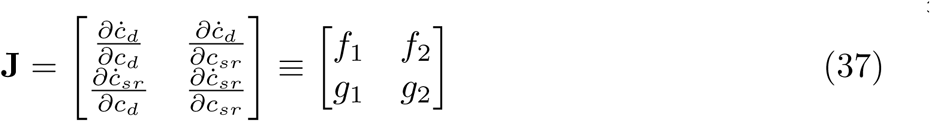

A fixed point will be stable provided *f*_1_ + *g*_2_ *<* 0. When *f*_1_*g*_2_ − *f*_2_*g*_1_ *>* 0 the stable fixed point corresponds to a node and if *f*_1_*g*_2_ – *f*_2_*g*_1_ *<* 0 to a stable spiral. We can use this to relate the slope of the nullclines to the stability of the upper fixed point. Let us denote *α* ≡ *ċ*_*d*_ and *β* ≡ *ċ*_*sr*_ the time derivatives of the independent variables.

At large values of the total concentration (see Fig. 13a, with 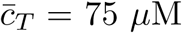), the slope of the black nullcline (*α* = 0) at the fixed point is positive, while the slope of the orange nullcline (*β* = 0) is negative. Then, increasing *c*_*d*_ at constant *c*_*sr*_, *α* goes from being positive to negative. This means that *f*_1_ ≡ *∂α*/*∂c*_*d*_ *<* 0. Using the same argument, it is easy to check that also *g*_2_ ≡ *∂β*/*∂c*_*sr*_ *<* 0, and therefore the fixed point is stable. At lower values of the total concentration (see, for instance, Fig. 13a, for 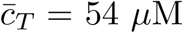) the slope of the black nullcline becomes negative. Thus, in this case, while *g*_2_ is still negative, *f*_1_ becomes positive, and it is not possible to determine the stability of the fixed point. It will depend on the speed of rate of *c*_*d*_ and *c*_*sr*_ close to the fixed point. If the dynamics of *c*_*d*_ is faster, then one expects this point to be unstable, if it is *c*_*sr*_ that varies fast, then stable. One would then expect that buffers that change the dynamics either in the cytosol or in the SR would effect the stability of the fixed point and, therefore, the range of existence of the limit cycle.

**Fig 13.**
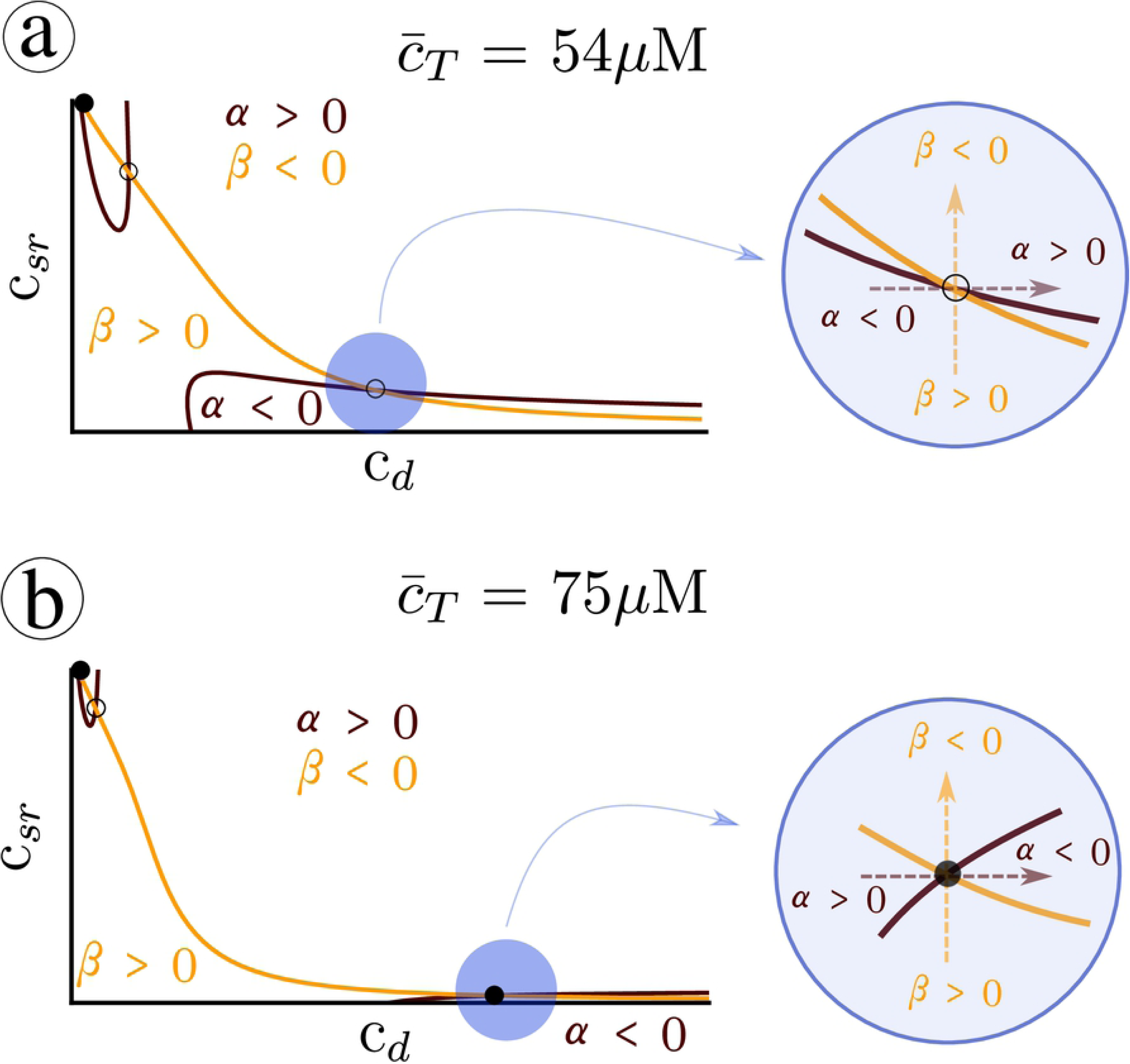
Structure of the nullclines at two different values of a) 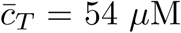, b) 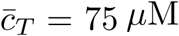. A black line corresponds to the nullcline *ċ*_*d*_ = 0, while the orange line indicates *ċ*_*sr*_ = 0. The functions *f*_1_ and *g*_2_ are the elements of the diagonal of the Jacobian matrix defined as 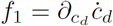 and 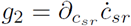.

### Robustness of the results. Fast dyadic calcium dynamics

It is useful to check that the basic points of our discussion hold when different possible approximations are applied to the minimal model. Namely, if the time scale of RyR opening is not as fast as the time scale of calcium diffusion near the dyadic space we should analyze the fast dyadic calcium approximation and not the fast RyR opening approximation to obtain information from the nullcline analysis. To this end, we have performed simulations of Eqs (21)-(22) at different values of the total calcium concentration 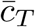 (Fig. 14) and test that we find the same basic behavior: at low values of 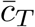 the system remains in a low concentration steady state (Fig. 15), but oscillations appear for a range of 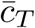, up to a limit where an upper state becomes stable.

**Fig 14.**
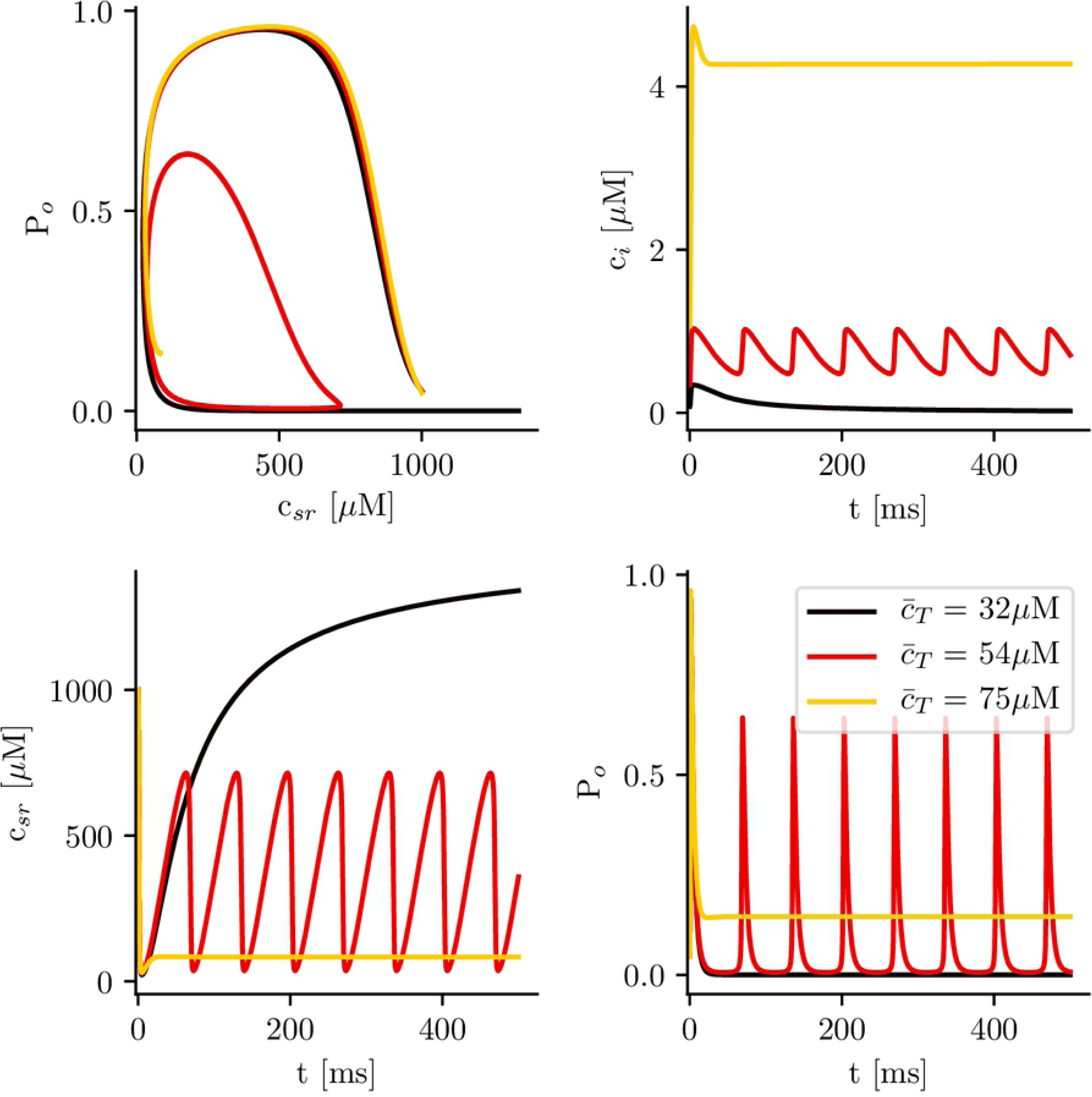
Solutions as a function of total calcium concentration 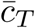, with calsequestrin concentration set to zero when the fast dyadic approximation is used. We obtain the same type of structure as expected.The system can be in a monostable state, which is excitatory (low load), in an oscillatory state (intermediate load), or in a bistable state (high loads), where it usually ends in a state of open RyR and depleted SR calcium concentration.

**Fig 15.**
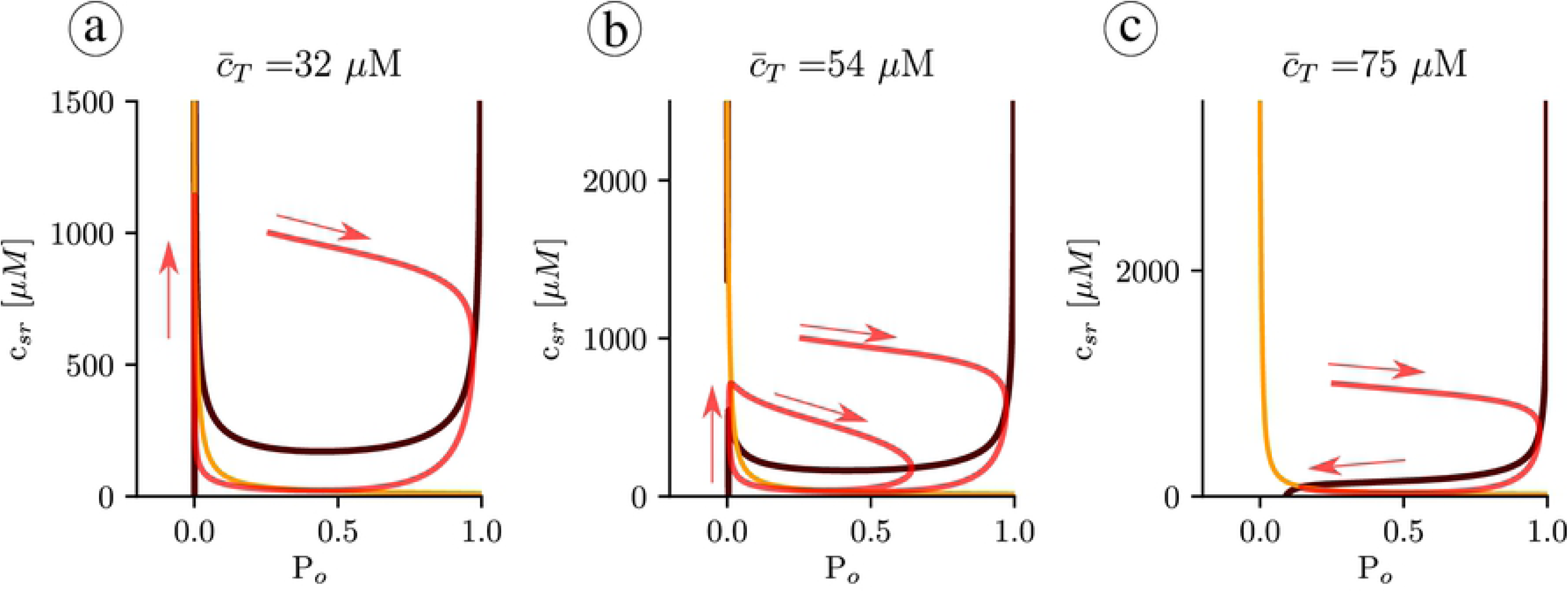
Structure of the nullclines at different values of 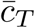 indicated in the title of each panel. The black line indicates the nullcline 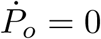, while the orange line corresponds to *ċ*_*sr*_ = 0. The red curve is a trajectory with a direction indicated with the red arrows.

More importantly, the basic structure of the nullclines determines the possible solutions and again, oscillations appear when pinch off is produced (Fig. 14). The fixed points in this case are the same as in Fig. 7, since they correspond to fixed points of Eqs. (12)-(14). However, different slaving conditions may change the stability of the fixed points, that now are analyzed in the plane (*c*_*sr*_, *P*_*o*_). Similarly to what we found in the previous analysis, calculating the stability, we find that the intermediate state is always unstable while the upper branch is stable above certain value of 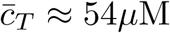. Both states appear at around 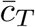, indicating the robustness of our analysis.

## Discussion and conclusion

Calcium oscillations play an important role in cardiac cells, from the regulation of growth in human cardiac progenitor cells [34], to the control of the pacemaker rhythm in both early embryonic heart cells [35, 36] and in sinoatrial nodal pacemaker cells (SANCs) [37–39]. Spontaneous calcium releases or oscillations are also related to the appearance of early or delayed-afterdepolarizations [40], which may give rise to reentrant waves and arrhythmic rhythms, such as tachycardia or fibrillation. Calcium oscillations have been observed under conditions of high cytosolic calcium concentration [23] or SR calcium overload. High levels of cytosolic calcium affect the opening probability of the RyR, which may result in oscillations or in a permanently open state [23, 41]. Calcium overload can be obtained, for instance, by inhibition of the Na^+^-K^+^ pump current *I*_*NaK*_, that results in [Na^+^]_*i*_ overload. The consequent build-up of [Na^+^]_*i*_ reduces the effectiveness of the Na^+^-Ca^2+^ exchanger at removing calcium from the cell and intracellular calcium concentrations become elevated. A similar effect is observed in models of hypercalcemia [42]. The effect of elevated [Na^+^]_*i*_ has been studied in computational models, finding calcium oscillations [43], that, depending on the model, appear via a supercritical Hopf [24] or a homoclinic bifurcation [25, 26].

We have shown, using a full subcellular model, that under global calcium overload, SR oscillations appear, leading finally to a state with permanently open RyR and depleted SR. In these simulations, oscillations give rise to periodic calcium waves, that propagate along the myocyte. To obtain a better understanding of the origin and parameter dependence of these transitions, we have studied them in a simplified model of the calcium dynamics. Despite not including all the physiological details (or because of that), minimal models are often useful to gain a better understanding of the origin of complex calcium rhythms [44], as oscillations [45]. We have thus analyzed the different transitions within a minimal model, that takes into account the RyR dynamics, as well as fluxes among different compartments. This allows us to explain the origin of oscillations using a nullcline analysis, as well as to give analytic expressions for the different transitions. One interesting conclusion is that buffers affect heavily the dynamics. The effect of buffers on oscillations has been studied previously [32], and experimentally a change in CSQ levels has been observed to alter the range of values of cytosolic calcium at which oscillations are observed [23]. Here we find that an increase in the levels of CSQ prevents altogether the oscillations, obtaining a direct transition to an open state, that also occurs at larger values of total calcium content. We have shown that this picture is robust and independent on which is faster, whether the opening of the RyR2 or the diffusion of calcium near it.

The fact that oscillations appear both in a fully detailed stochastic model and in a simple deterministic model seems another proof of the robustness of the behavior. This robustness is not unexpected since it has already been observed in other calcium signaling processes, such as calcium oscillations in hepatocytes [46, 47]. Nevertheless, while the minimal deterministic model presents a sudden drop in SR calcium content at the onset of oscillations, the subcellular stochastic model presents a gradual decrease in the level of basal SR, in full agreement with the experimental results obtained by Stevens et al [23]. Thus, the comparative analysis of our results seems to suggest that the origin of oscillations in experiments arises from the smearing out of a sharp transition to the oscillatory state, due to the stochasticity of the RyR in an extended system.

Finally, in this paper, we consider a cell which has achieved calcium balance where intrusion and extrusion match, and have neglected calcium fluxes across the cell membrane to focus on the internal calcium dynamics in order to decouple both processes. Under normal pacing, extracellular calcium fluxes typically represent about 10-20% of the total calcium fluxes, so it is not unreasonable to consider that the total calcium content remains constant once at a steady state. Under these conditions, cytosolic and SR calcium concentrations are not independent but linked, and a clear control parameter is the total calcium concentration. Here we show that it uniquely determines the state of the system. Of course, in the presence of transmembrane fluxes, calcium oscillations or waves, or a permanently open RyR, would result in an extrusion of calcium out of the cell and an eventual decrease in the total calcium load of the cell, that would transition back to the quiescent state (in the absence of external pacing). It seems interesting to study in the future the effect of oscillations and waves in the action potential, as well as a paced cell at different periods and the interaction with the pacing period. Observing how the time scale related to oscillations interacts with the time scale needed to extrude the calcium in the cytosol if the open (upper state) is reached may lead to new interesting phenomena.

## Acknowledgments

Financial support was provided by the Fundació La Marató de TV3, under grant number 20151110, and from the Spanish Ministerio de Economía, Industria y Competitividad under grant number SAF2017-88019-C3-2-R. Support for YS was provided by NHLBI grant R01-HL119095. We want to thank I.R.Cantalapiedra, A.Peñaranda and L.Hove-Madsen from fruitful discussions.

## A Full calcium model of a CaRU

In this appendix, we show how to derive the simplified model in Eqs. (12)-(14) from a detailed deterministic model of calcium handling. We consider cytosolic and luminal spaces, each separated into different compartments, i.e, dyadic and cytosolic by the one side, and junctional and network SR, by the other. We also consider the effect of two buffers in the cytosol, TnC and SR, and Calsequestrin (CSQ) in the SR. The effect of the buffers is actually very relevant, as they change the structure of possible solutions of the system.

With this, the dynamics can be described by the following set of deterministic equations for the calcium concentration at the different compartments

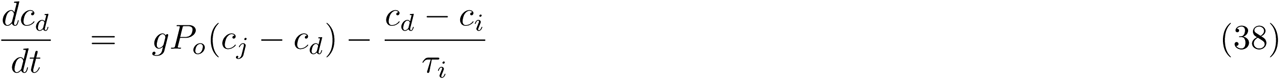

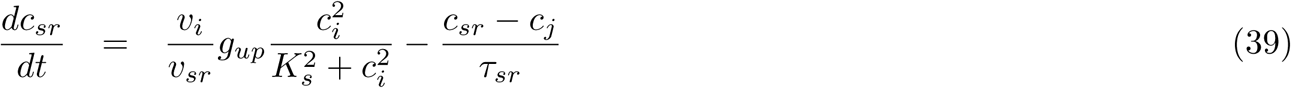

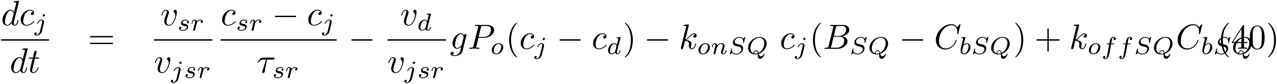

where *c*_*d*_, *c*_*j*_, *c*_*sr*_, *c*_*i*_ stand for the concentration in dyadic space, free luminal, SR network and cytosol, *P*_*o*_ is the fraction of RyRs in the open state, *τ*_*i*_ and *τ*_*sr*_ are the diffusion time constants out of the dyadic space and SR network, *v*_*i*_, *v*_*d*_, *v*_*jsr*_ and *v*_*sr*_ are the volumes associated with each compartment, *g*_*up*_ is the strength of SERCA pump, *K*_*s*_ is the concentration at which SERCA closes and *g* is the strength of the release current. The dynamics of the buffers are given by linear reactions with the following set of ODEs

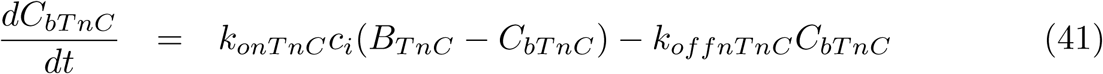

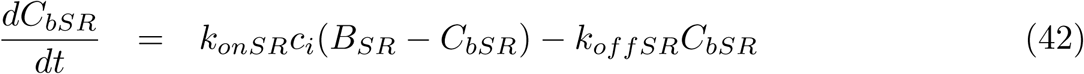

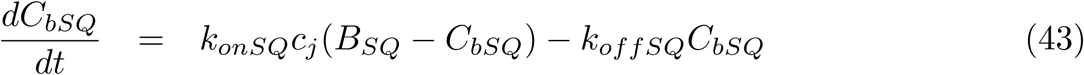

Since the total amount of calcium has just a variation of about by a 5% or 10% over a calcium cycle, we assume that the total calcium concentration, 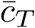, is fixed. In this way, one does not have to solve a differential equation for the calcium concentration in the cytosol. Rather, it is derived from the algebraic equation:

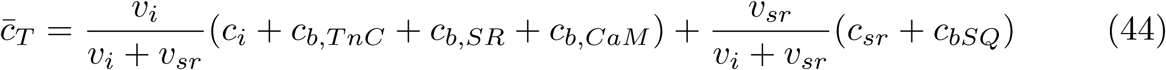

Homeostatic behavior of the cell will eventually load the system more or less, increasing or decreasing 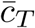.

Gating of the RyR can been described by Markov models that describe the transitions among different conformations of the channel. Thus, for the dynamics of the RyR we consider a phenomenological four state model [48].

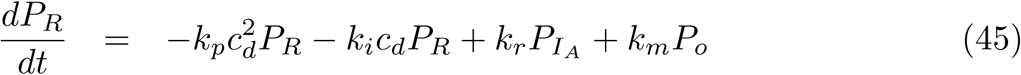

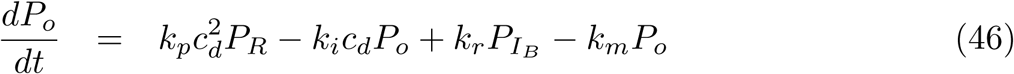

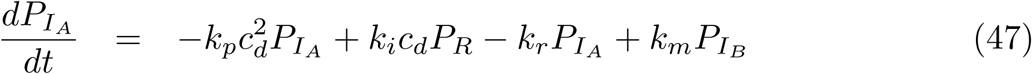

with the last equation given by the condition that the sum of probabilities is equal to 1

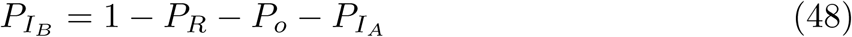

*P*_*R*_ and *P*_*o*_ are the ratios of local RyR in the recovered and open states. 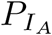 and 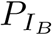 stand for the terminated states.

To make the model treatable we assume several hypotheses.

1. We consider rapid equilibrium of *c*_*j*_ (*ċ*_*j*_ = 0). Thus

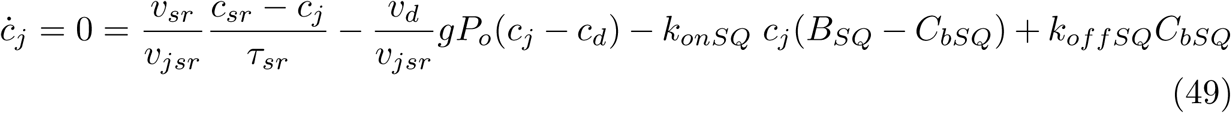

then

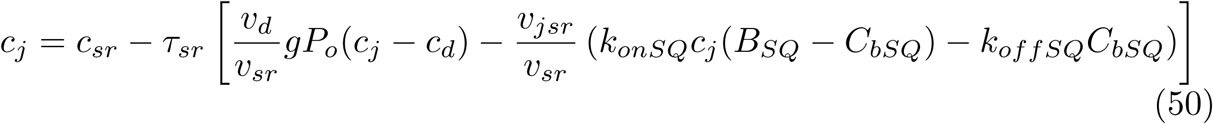
2. At first order, we approximate *c*_*j*_ as *c*_*sr*_ in the right hand side of Eq. (50). This approximation is valid when 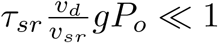 and 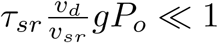. Then, the ODE for *c*_*sr*_ in Eq. (40) reads as

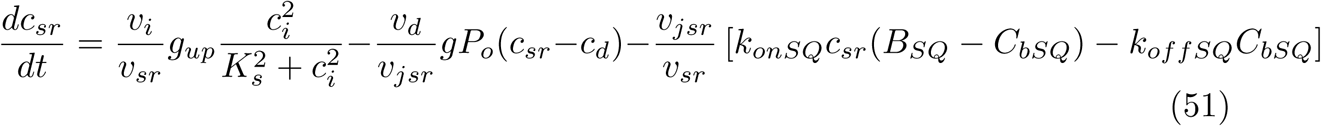

where we have eliminated the dependence with *c*_*j*_.
3. We apply the rapid buffer approximation in the SR because CSQ is very fast [29, 30]. The derivation of *c*_*sr*_ in terms of 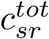 have been already shown in the main text (see Eqs. 7-10).
4. We assume that all the buffers in the cytosol are in equilibrium. Then

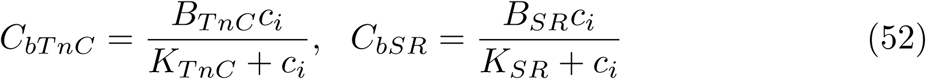
5. Besides, we combine both TnC and SR buffers in only one buffer with a total concentration of *B*_*b*_ and an affinity of *K*_*b*_, which will be

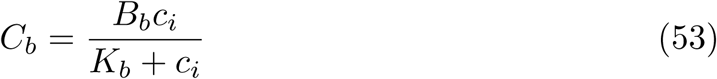
6. Finally, we consider that the inactivated states of the RyRs do not play a role in the mechanisms that produce oscillations. For that reason, we reduce the four state model to a two state model

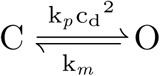

where *k*_*m*_ sets the mean open time *τ*_*rec*_ = 1/*k*_*m*_ of a RyR while *k*_*p*_ gives the open probability. Then, the ODE for the open probability *P*_*o*_ is

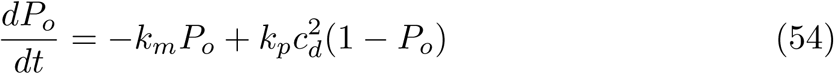

The parameters of these equations are taken from the literature where, except for those of the RyR, are well documented. The ratio of SR to cytosol in the cell is roughly 1 to 20-30. The order of magnitude of the volume of the cleft where calcium is released is around 10^−3^ *µm*^3^. SERCA is roughly activated at around 2-5 *µM* and closed at around 15-20 *µM* with the number of buffers relevant to absorption around 50-100 *µM*, taking an average affinity of around 0.5 *µM*. The whole set of parameters is given in Table 1.

## References

1. Mendis S, Puska P, Norrving B, Organization WH, et al. Global atlas on cardiovascular disease prevention and control. Geneva: World Health Organization; 2011.

2. Huikuri HV, Castellanos A, Myerburg RJ. Sudden death due to cardiac arrhythmias. New England Journal of Medicine. 2001;345(20):1473–1482.

3. Qu Z, Weiss JN. Mechanisms of ventricular arrhythmias: from molecular fluctuations to electrical turbulence. Annual review of physiology. 2015;77:29–55.

4. Alonso S, Bär M, Echebarria B. Nonlinear physics of electrical wave propagation in the heart: a review. Reports on Progress in Physics. 2016;79(9):096601.

5. Song Z, Qu Z, Karma A. Stochastic initiation and termination of calcium-mediated triggered activity in cardiac myocytes. Proceedings of the National Academy of Sciences. 2017;114(3):E270–E279.

6. Colman MA. Arrhythmia mechanisms and spontaneous calcium release: Bi-directional coupling between re-entrant and focal excitation. PLoS computational biology. 2019;15(8).

7. Hoeker GS, Katra RP, Wilson LD, Plummer BN, Laurita KR. Spontaneous calcium release in tissue from the failing canine heart. American Journal of Physiology-Heart and Circulatory Physiology. 2009;297(4):H1235–H1242.

8. Pogwizd SM, McKenzie JP, Cain ME. Mechanisms underlying spontaneous and induced ventricular arrhythmias in patients with idiopathic dilated cardiomyopathy. Circulation. 1998;98(22):2404–2414.

9. Zhou S, Chang CM, Wu TJ, Miyauchi Y, Okuyama Y, Park AM, et al. Nonreentrant focal activations in pulmonary veins in canine model of sustained atrial fibrillation. American Journal of Physiology-Heart and Circulatory Physiology. 2002;283(3):H1244–H1252.

10. Orchard C, Eisner D, Allen D. Oscillations of intracellular Ca2+ in mammalian cardiac muscle. Nature. 1983;304(5928):735.

11. Kort AA, Lakatta EG. Calcium-dependent mechanical oscillations occur spontaneously in unstimulated mammalian cardiac tissues. Circulation research. 1984;54(4):396–404.

12. Cheng H, Lederer M, Lederer W, Cannell M. Calcium sparks and [Ca2+]i waves in cardiac myocytes. American Journal of Physiology-Cell Physiology. 1996;270(1):C148–C159.

13. Marchant JS, Parker I. Role of elementary Ca 2+ puffs in generating repetitive Ca 2+ oscillations. The EMBO Journal. 2001;20(1-2):65–76.

14. Uhlén P, Fritz N. Biochemistry of calcium oscillations. Biochemical and biophysical research communications. 2010;396(1):28–32.

15. Dupont G, Combettes L, Bird GS, Putney JW. Calcium oscillations. Cold Spring Harbor perspectives in biology. 2011;3(3):a004226.

16. Plummer BN, Cutler MJ, Wan X, Laurita KR. Spontaneous calcium oscillations during diastole in the whole heart: the influence of ryanodine reception function and gap junction coupling. American Journal of Physiology-Heart and Circulatory Physiology. 2011;300(5):H1822–H1828.

17. Katra RP, Oya T, Hoeker GS, Laurita KR. Ryanodine receptor dysfunction and triggered activity in the heart. American Journal of Physiology-Heart and Circulatory Physiology. 2007;292(5):H2144–H2151.

18. Priori SG, Napolitano C, Tiso N, Memmi M, Vignati G, Bloise R, et al. Mutations in the cardiac ryanodine receptor gene (hRyR2) underlie catecholaminergic polymorphic ventricular tachycardia. Circulation. 2001;103(2):196–200.

19. Lou Q, Belevych AE, Radwański PB, Liu B, Kalyanasundaram A, Knollmann BC, et al. Alternating membrane potential/calcium interplay underlies repetitive focal activity in a genetic model of calcium-dependent atrial arrhythmias. The Journal of physiology. 2015;593(6):1443–1458.

20. Van Petegem F. Ryanodine receptors: structure and function. Journal of Biological Chemistry. 2012;287(38):31624–31632.

21. Xie LH, Weiss JN. Arrhythmogenic consequences of intracellular calcium waves. American Journal of Physiology-Heart and Circulatory Physiology. 2009;297(3):H997–H1002.

22. Shiferaw Y, Aistrup GL, Wasserstrom JA. Intracellular Ca2+ waves, afterdepolarizations, and triggered arrhythmias; 2012; Cardiovascular Research, 95(3), 265.

23. Stevens SC, Terentyev D, Kalyanasundaram A, Periasamy M, Györke S. Intra-sarcoplasmic reticulum Ca2+ oscillations are driven by dynamic regulation of ryanodine receptor function by luminal Ca2+ in cardiomyocytes. The Journal of physiology. 2009;587(20):4863–4872.

24. Varghese A, Winslow RL. Dynamics of the calcium subsystem in cardiac Purkinje fibers. Physica D: Nonlinear Phenomena. 1993;68(3-4):364–386.

25. Winslow RL, Varghese A, Noble D, Adlakha C, Hoythya A. Generation and propagation of ectopic beats induced by spatially localized Na–K pump inhibition in atrial network models. Proceedings of the Royal Society of London Series B: Biological Sciences. 1993;254(1339):55–61.

26. Benson AP, Holden AV. Calcium oscillations and ectopic beats in virtual ventricular myocytes and tissues: bifurcations, autorhythmicity and propagation. In: International Workshop on Functional Imaging and Modeling of the Heart. Springer; 2005. p. 304–313.

27. Marchena M, Echebarria B. Computational model of calcium signaling in cardiac atrial cells at the submicron scale. Frontiers in physiology. 2018;9:1760.

28. Song Z, Karma A, Weiss JN, Qu Z. Long-lasting sparks: Multi-metastability and release competition in the calcium release unit network. PLoS computational biology. 2016;12(1):e1004671.

29. Robertson S, Johnson JD, Potter J. The time-course of Ca2+ exchange with calmodulin, troponin, parvalbumin, and myosin in response to transient increases in Ca2+. Biophysical journal. 1981;34(3):559–569.

30. Cannell M, Allen D. Model of calcium movements during activation in the sarcomere of frog skeletal muscle. Biophysical Journal. 1984;45(5):913–925.

31. Donoso P, Prieto H, Hidalgo C. Luminal calcium regulates calcium release in triads isolated from frog and rabbit skeletal muscle. Biophysical journal. 1995;68(2):507–515.

32. Wagner J, Keizer J. Effects of rapid buffers on Ca2+ diffusion and Ca2+ oscillations. Biophysical Journal. 1994;67(1):447–456.

33. Ermentrout B. Simulating, analyzing, and animating dynamical systems: a guide to XPPAUT for researchers and students. vol. 14. Siam; 2002.

34. Ferreira-Martins J, Rondon-Clavo C, Tugal D, Korn JA, Rizzi R, Padin-Iruegas ME, et al. Spontaneous calcium oscillations regulate human cardiac progenitor cell growth. Circulation research. 2009;105(8):764–774.

35. Viatchenko-Karpinski S, Fleischmann B, Liu Q, Sauer H, Gryshchenko O, Ji G, et al. Intracellular Ca2+ oscillations drive spontaneous contractions in cardiomyocytes during early development. Proceedings of the National Academy of Sciences. 1999;96(14):8259–8264.

36. Sasse P, Zhang J, Cleemann L, Morad M, Hescheler J, Fleischmann BK. Intracellular Ca 2+ oscillations, a potential pacemaking mechanism in early embryonic heart cells. The Journal of general physiology. 2007;130(2):133–144.

37. Lakatta EG, Maltsev VA, Vinogradova TM. A coupled SYSTEM of intracellular Ca2+ clocks and surface membrane voltage clocks controls the timekeeping mechanism of the heart’s pacemaker. Circulation research. 2010;106(4):659–673.

38. Vinogradova TM, Zhou YY, Maltsev V, Lyashkov A, Stern M, Lakatta EG. Rhythmic ryanodine receptor Ca2+ releases during diastolic depolarization of sinoatrial pacemaker cells do not require membrane depolarization. Circulation research. 2004;94(6):802–809.

39. Vinogradova TM, Lyashkov AE, Zhu W, Ruknudin AM, Sirenko S, Yang D, et al. High basal protein kinase A–dependent phosphorylation drives rhythmic internal Ca2+ store oscillations and spontaneous beating of cardiac pacemaker cells. Circulation research. 2006;98(4):505–514.

40. Spencer CI, Sham JS. Effects of Na+/Ca2+ exchange induced by SR Ca2+ release on action potentials and afterdepolarizations in guinea pig ventricular myocytes. American Journal of Physiology-Heart and Circulatory Physiology. 2003;285(6):H2552–H2562.

41. Saftenku E, Williams AJ, Sitsapesan R. Markovian models of low and high activity levels of cardiac ryanodine receptors. Biophysical journal. 2001;80(6):2727–2741.

42. Alvarez-Lacalle E, Peñaranda A, Cantalapiedra IR, Hove-Madsen L, Echebarria B. Effect of RyR2 Refractoriness and Hypercalcemia on Calcium Overload, Spontaneous Release, and Calcium Alternans. Computing in Cardiology 2013; 40:683–686.

43. Varghese A, Winslow RL. Dynamics of abnormal pacemaking activity in cardiac Purkinje fibers. Journal of theoretical biology. 1994;168(4):407–420.

44. Cantalapiedra IR, Alvarez-Lacalle E, Peñaranda A, Echebarria B. Minimal model for calcium alternans due to SR release refractoriness. Chaos: An Interdisciplinary Journal of Nonlinear Science. 2017;27(9):093928.

45. Sneyd J, Han JM, Wang L, Chen J, Yang X, Tanimura A, et al. On the dynamical structure of calcium oscillations. Proceedings of the National Academy of Sciences. 2017;114(7):1456–1461.

46. Kummer U, Krajnc B, Pahle J, Green AK, Dixon CJ, Marhl M. Transition from stochastic to deterministic behavior in calcium oscillations. Biophysical journal. 2005;89(3):1603–1611.

47. Dupont, G, Croisier, H. Spatiotemporal organization of Ca2+ dynamics: A modeling-based approach. HFSP journal. 2010;4(2): 43–51.

48. Stern MD, Song LS, Cheng H, Sham JS, Yang HT, Boheler KR, et al. Local control models of cardiac excitation–contraction coupling. The Journal of general physiology. 1999;113(3):469–489.

